# *In vitro* pharmacokinetics and pharmacodynamics of the diarylquinoline TBAJ-587 and its metabolites against *Mycobacterium tuberculosis*

**DOI:** 10.1101/2025.11.05.686716

**Authors:** Diana Angelica Aguilar-Ayala, Marie Sylvianne Rabodoarivelo, Maxime R. Eveque-Mourroux, Albin AM Leding, Lindsay Sonnekalb, Ana Picó Marco, Nicolas Willand, Natalya Serbina, Ulrika SH Simonsson, Ainhoa Lucía, Santiago Ramón-García, the ERA4TB consortium

**Affiliations:** Department of Microbiology, Pediatrics, Radiology and Public Health, Faculty of Medicine, University of Zaragoza, Spain; Univ. Lille, Inserm, Institut Pasteur Lille, U1177 - Drugs and Molecules for living Systems, F-59000 Lille, France; Dept. of Pharmaceutical Biosciences, Uppsala University, Uppsala, Sweden; Molecular and Experimental Mycobacteriology, Research Center Borstel Leibniz Lung Center, Parkallee 1-40, 23845, Borstel, Germany; The Global Alliance for TB Drug Development, New York, NY, United States; Spanish Network for Research on Respiratory Diseases (CIBERES), Carlos III Health Institute, Madrid, Spain; Research & Development Agency of Aragón Foundation (Fundación ARAID), Zaragoza, Spain

**Keywords:** tuberculosis, TBAJ-587, pharmacokinetics/pharmacodynamics, time-kill assay

## Abstract

The first-in-class diarylquinoline (DARQ) bedaquiline (BDQ) is in the medicines list for drug-resistant tuberculosis. TBAJ-587 is a next-generation DARQ with improved anti-*Mycobacterium tuberculosis* (*Mtb*) activity and reduced cardiac repolarization abnormalities.

Methods. The *in-vitro* efficacy of TBAJ-587 and its main metabolites (M2, M3 and M12) was analyzed under standard (ST) growth conditions, with cholesterol (CHO), or fatty acids (FA) as alternative carbon sources. Minimal inhibitory concentration (MIC) assays and time-kill assays (TKA) linked to drug measurements in bacterial samples were performed to allow correlation of pharmacodynamics (PD) with actual pharmacokinetics (PK).

The most active compounds, TBAJ-587 and its M3 metabolite, exhibited broth media and concentration dependent efficacy showing a bactericidal effect at ≥5x MIC. Bacterial cultures treated with 1x MIC and 2x MIC of TBAJ-587 resumed growth after 28 days and displayed moderate increased MIC values compared to untreated conditions, which were linked to new variants of BDQ resistance mutations in the *atpE*, *atpB,* and *Rv0678* genes.

This study revealed TBAJ-587 and metabolites bind to polystyrene plastic-ware, the most commonly used material in antimicrobial research, being the effective unbound drug concentration dependent on the media composition. PKPD analyses determined that *Mtb* was killed with lower exposures of TBAJ-587 and M3 than expected in ST and FA broth, suggesting previously underestimated potency in these media.

This is the first *in-vitro* study to precisely link compound activities (PD) to their effective concentration (PK) over time in *Mtb* cultures, providing improved longitudinal data to feed models for translational research.

## INTRODUCTION

Bedaquiline (BDQ) is the first diarylquinoline (DARQ) approved by the World Health Organization (WHO) for use in combinatorial regimens against multi-drug resistant tuberculosis (MDR-TB). It is equally active against drug susceptible *Mycobacterium tuberculosis* (*Mtb*) strains and *Mtb* isolates resistant to isoniazid, rifampin, streptomycin, ethambutol, pyrazinamide, and moxifloxacin ^1^. BDQ specifically inhibits the mycobacterial ATP synthase minimizing the damage to healthy microbiota and reducing the chances of drug resistance development in non-mycobacterial populations. DARQs are the first anti-tuberculosis (TB) drugs with a distinct mechanism of action from other clinically approved treatments. This difference makes DARQs particularly effective when used in drug combinations, as they can together target different bacterial metabolic pathways. This approach increases the likelihood of eliminating recalcitrant bacterial subpopulations that may survive treatment with other anti-TB drugs. Despite its benefits, BDQ has cardiovascular side effects. Its major metabolite, M2, mainly produced through BDQ N-demethylation by the CYP3A4 in the liver, is less effective than BDQ and accumulates intracellularly at higher levels ^2,3^. Both BDQ and M2, block repolarization of hERG channels of cardiac cells, with M2 causing a stronger phospholipidosis than BDQ ^4,5^. Due to their high lipophilicity (BDQ-ClogP = 7.25, and M2-ClogP = 6.59), both compounds are prone to excessive absorption in fatty tissues and non-target sites. These drugs have a long terminal half-life of 4 - 5 months in humans; however, effective half-life is approximately 24 hours after just two weeks of daily dosing ^5^. These properties increase the risk of toxicity even after treatment completion^6^. Since the introduction of BDQ to the clinic, resistance-associated variants (RAVs) in the *atpE* and *Rv0678* genes of *Mtb* have been linked to reduced treatment efficacy. RAVs in *atpE* are rare in clinical strains, while Rv0678 RAVs are more frequent in baseline isolates of MDR-TB patients, resulting in a 2–8-fold increase in MIC and low-level resistance to BDQ ^7,8^. These findings emphasize the need for next-generation DARQs with improve potency to overcome the challenge of emerging resistance while maintaining acceptable safety margins.

Over the past decades, analogues of BDQ have been developed and investigated ^9–14^. The replacement of the naphthalene C unit by a pyridyl one led to the development of TBAJ-587, a DARQ retaining the mechanism of action against the mycobacterial ATP synthase, exhibiting lower lipophilicity with enhanced antibacterial activity against *Mtb* in *in-vitro* and mice assays. TBAJ-587 also maintains activity against wild type and Rv0678 mutant strains ^10,15,16^, with a superior cardiovascular safety profile than BDQ ^17^. In addition, TBAJ-587 performs better than BDQ in rabbit necrotic granuloma models by reaching bactericidal concentrations in the caseum cavities faster, thereby minimizing the chances of resistance development, in contrast to the slower diffusion kinetics of BDQ ^18^.

A phase 1 single and multiple ascending dose clinical trial of TBAJ-587 (ClinicalTrials.gov number, NCT04890535) was recently completed within the ERA4TB consortium (https://era4tb.org/); this endorsement enables extensive research within ERA4TB facilitating a robust integration of *in-vitro* and *in vivo* studies to refine the selection of effective doses for translational applications. Here, we implemented for the first time a multimodal *in-vitro* pharmacokinetic/pharmacodynamic (PKPD) approach to characterize the activity of the TBAJ-587 and its main metabolites (M2, M3 and M12) against *Mtb* H37Rv. Our strategy determined the link between the actual effective concentration in static conditions over time in different carbon sources and the anti-mycobacterial activity of the compounds demonstrating a comprehensive PKPD relationship. Our *in-vitro* assays are a key addition to the drug development process to better characterize PKPD parameters, which will feed PKPD models for dose optimization clinical translation studies.

## METHODS

### Bacterial strain, culture conditions and drugs

*Mtb* H37Rv (GenBank ID: NC_000962.3) was used for all assays. Bacterial stocks were generated in Difco^TM^ Middlebrook 7H9 with 0.5% glycerol, 10% BBL^TM^ Middlebrook OADC and 0.25% tyloxapol, as described previously ^19^, and stored at - 80°C. Three stock tubes are quantified on the day of stock preparation and 14 days after storage for CFU enumeration. Characterized vials were thawed each time for every assay in polystyrene flasks in the correspondent media broth, and adjusted to 10^4^ CFU/mL for three days recovery at 37°C, before drug compounds were added into bacterial cultures. Eight media broth were used: (i) *standard* (ST): 7H9 + 10% OADC + 0.5% glycerol; (ii) *cholesterol* (CHO): 7H9 + 0.5% BSA + 0.085% NaCl + 0.05% tyloxapol + 0.004% cholesterol; (iii) *fatty acids* (FA): 7H9 + 0.5% BSA + 0.085% NaCl + 0.001% palmitic acid + 0.001% stearic acid + 0.001% oleic acid; (iv) ST broth supplemented with *tyloxapol* (for comparison with the CHO broth): 7H9 + 10% OADC + 0.5% glycerol + 0.05% tyloxapol; (v) *glucose* broth: 7H9 + 0.5% BSA + 0.085% NaCl + 1% glucose; (vi) *pyruvate*: 7H9 + 0.5% BSA + 0.085% NaCl + 20 mM pyruvate; (vii) *acetate*: 7H9 + 0.5% BSA + 0.085% NaCl + 10 mM sodium acetate and; (viii) *butyrate*: 7H9 + 0.5% BSA + 0.085% NaCl + 5mM sodium butyrate. Lyophilized DARQ compounds were provided by TB-Alliance. BDQ fumarate, TBAJ-587 fumarate and TBAJ-587 M2, TBAJ-587 M3, and TBAJ-587 M12 metabolites were solved in DMSO at 10 mg/mL or 1 mg/mL in glass vials covered with aluminum foil, then stored at -20°C until their use in this experiment. Polypropylene material was avoided ^20^, whenever further dilutions of the master stocks were needed, glass tubes and polystyrene plastic ware was preferred. Only polypropylene tips could not be substituted. Assays were carried out either in polystyrene microplates or flasks. Moxifloxacin (MXF, European Pharmacopoeia Reference Standard, Y0000703, Batch 3.0), and linezolid (LZD, SIGMA-ALDRICH, USP Reference Standard, 1367561, batch R072F0) were used as internal controls. Lyophilized MXF and LZN were solved in distilled water at 1 mg/mL, filtered and stored at -20°C until their use in this experiment.

### Minimal inhibition concentration (MIC) determinations

MIC assays of DARQs were performed against *Mtb* H37Rv from characterized stocks of the *Mtb* H37Rv wild type strain using the eight media broths mentioned above. Tests using MXF, LZD, untreated and sterility controls were included. MIC values were determined through the broth micro-dilution resazurin microtiter assay (REMA) method^21^. Bacterial pre-inocula at 10^4^ CFU/mL were incubated for two days in static conditions at 37°C without CO_2_ before being added to the 96-well-microplates containing a 2-fold serial dilution concentration range. At least three technical replicates were carried out for each determination. Cultures were incubated for six days in the presence of the compounds, before addition of 30 μL of 0.01% diluted resazurin. Growth was evaluated by reading fluorescence (Excitation: 530 nm / 25 nm, Emission: 590 nm / 20 nm) after 24 h of resazurin addition using a Synergy HT Biotek® microplate reader (Synergy HTX BioTek Gen5 v 2.06 software). Reported MIC values correspond to MIC_80_, which is the lowest drug concentration that inhibits ≥80% of bacterial growth.

### Dose response time kill assays

Time-kill assays (TKA) were performed for TBAJ-587 and its M2, M3, and M12 metabolites against *Mtb* H37Rv in ST, CHO, and FA broth media. Nine MIC-scaled concentrations were tested for each compound, including 1/20x, 1/5x, 1/2x, 1x, 2x, 5x, 20x, 100x, and 300x the MIC (Table S1). Scaled concentrations derived from MIC results in ST, CHO and FA conditions, MIC_TBAJ-587_ = 0.031 mg/L, MIC_M2_ = 0.4 mg/L, MIC_M3_ = 0.13 mg/L, and MIC_M12_ = 4.5 mg/L Untreated controls were also included for a total of 111 conditions. Thawed stocks of *Mtb* were adjusted to 10^4^ CFU/mL in ST, CHO or FA broth media in polystyrene tissue culture flasks (TPP® 25 cm^2^) and incubated for 3 days at 37°C in static conditions, without CO_2_, in order to allow for bacterial recovery up to ca. 10^5^ CFU/mL before compound addition. Cultures were then further incubated for 28 days at 37°C. Nine time-points were sampled for PD evaluation (CFU enumeration) and six time-points for PK measurements (drug quantification). Bacterial survival was measured by plating undiluted cultures and 10-fold serial dilutions (2.5 µL) on 7H10 + 10% OADC + 0.5% Glycerol + 0.1% tyloxapol agar plates on days 0, 1, 2, 4, 7, 10, 14, 21 and 28, as previously described ^19^. In parallel, 200 μL samples were stored in 96-well polystyrene microplates at -80°C on days 0, 2, 7, 14, 21, and 28 for drug quantification.

### Stability assays

Six TKA time-points for drug quantification, and four compounds (TBAJ-587, M2, M3 and M12), each at nine tested concentrations in three broth media (ST, CHO and FA) yielded a total of 648 samples kept at -80°C until evaluation. Samples were thawed and processed all at once, centrifuged, and bacterial pellets discarded. Decontamination was carried out by transferring supernatants into cold acetonitrile (AcN) in a 1:5 proportion, contained in polypropylene tubes. Samples were immediately stored at -80°C before being transferred into matrix tubes for LC-MS/MS analysis. A calibration curve containing each compound (TBAJ-587 and its metabolites) was established from 0 nM to 1000 nM in each media. Each point of the curve was performed in duplicate. For each sample point, proteins were precipitated with AcN and supernatants were transferred into matrix tubes for LC-MS/MS analysis.

LC-MS/MS analyses were performed using an UPLC Acquity I-Class system (Waters®) equipped with an Ethylene Bridge Hybrid C18 column (1,7 μm, 50 mm*2.1 mm) coupled online with a mass spectrometer instrument (Xevo TQ-D system, Waters®). The elution was performed with a 4 min gradient from 2% to 98% AcN with a flow rate set at 600 μL/min. The mass spectrometer equipped with an ESI source was operated in positive ion polarity with a specific MRM method for each compound of interest. Raw data were processed within the MassLynx software (version 4.1, Waters®).

To assess the effect of tyloxapol (contained in CHO broth) on TBAJ-587 compound stability, two TBAJ-587 concentrations were incubated at 37°C in 7H9 + 0.5% glycerol + 10% OADC media broth with and without tyloxapol (0.05%), without bacteria, over 28 days in polystyrene flasks. A low (0.16 mg/L) and a high concentration (1.6 mg/L) were tested and samples taken and quantified at 0, 1, 10 and 28 days after compound addition. Parallel conditions with propranolol (2 µM and 0.2 µm) were included as a drug control; two time-points were sampled for quantification at days 0 and 28 after compound addition. All drug measurements were performed in duplicates and verapamil was used as internal standard.

### PKPD data analysis

Bacterial survival at every time-point was transformed to relative bacterial burden to baseline, where baseline is the burden without treatment in each carbon source, and media broth was set as a batch effect. Graphical analysis was generated in RStudio (version 4.1.2) ^22^ and GraphPad software (version 8.0.2) ^23^. Exposures of drugs for each condition were derived as the area under the curve (AUC) calculated with ncappc version 0.3.0^24^, either assuming constant concentration over time (expected exposure) or calculated based on observed concentration over time (actual exposure). Relative bacterial burden to baseline was calculated according to **Equation 1**.

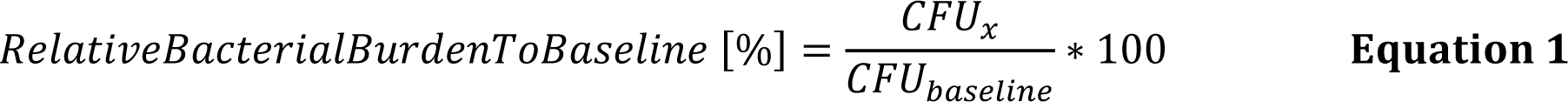

where CFU_x_ represent the observed readout, and CFU_baseline_ represents the corresponding median in the natural growth within the same medium at time point day 0 of the experiment.

### Whole Genome Sequencing

Libraries were made using Baym library preparation and sequenced on the Illumina NextSeq1000 platform. Whole genome sequencing analysis was performed using MTBseq pipeline set at low-frequency mode and genomes were aligned to the H37Rv reference genome (GenBank number NC_000962). Variant calling was customized by excluding all calls which did not fit the following criteria: were in a non-repetitive gene, were in a coding gene, had at least two reads in both forward and reverse orientation, had a frequency over 10%, and SNPs resulted in a missense mutation.

## RESULTS

### Anti-mycobacterial activity of TBAJ-587 and its main metabolites in eight different media

The MIC distribution for BDQ, TBAJ-587 and TBAJ-587 main metabolites in media with different sole carbon sources is captured in **Figure 1** and Table S2. Cholesterol, fatty acids, standard broth with tyloxapol, glucose, butyrate, pyruvate and acetate broth conditions were compared with standard cultures (ST). MIC values of TBAJ-587 ranged between 0.031 and 0.063 mg/L, which is no more than a 2-fold change across all eight-broth media. MIC values for BDQ shifted within a slightly higher and wider range than for TBAJ-587. MICs_BDQ_ in six carbon sources ranged between 0.063 and 0.25 mg/L, while the lowest MIC_BDQ_ was found in pyruvate (0.031 mg/L) and the highest in acetate broth (0.25 mg/L). MIC values for the M3 metabolite ranged from 0.063 to 0.125 mg/L in all broth media types except in acetate at 0.031 mg/L. A wider range of MIC values among carbon sources were found when using M2 and M12 metabolites: 0.2 – 3.2 mg/L and 1.125 – 9 mg/L, respectively, but MIC shifts were not specific to any individual carbon source. For all compounds, tyloxapol did not change the MIC more than 2-folds in the CHO broth and the ST plus tyloxapol broth in comparison with ST broth without tyloxapol.

**Figure 1.**
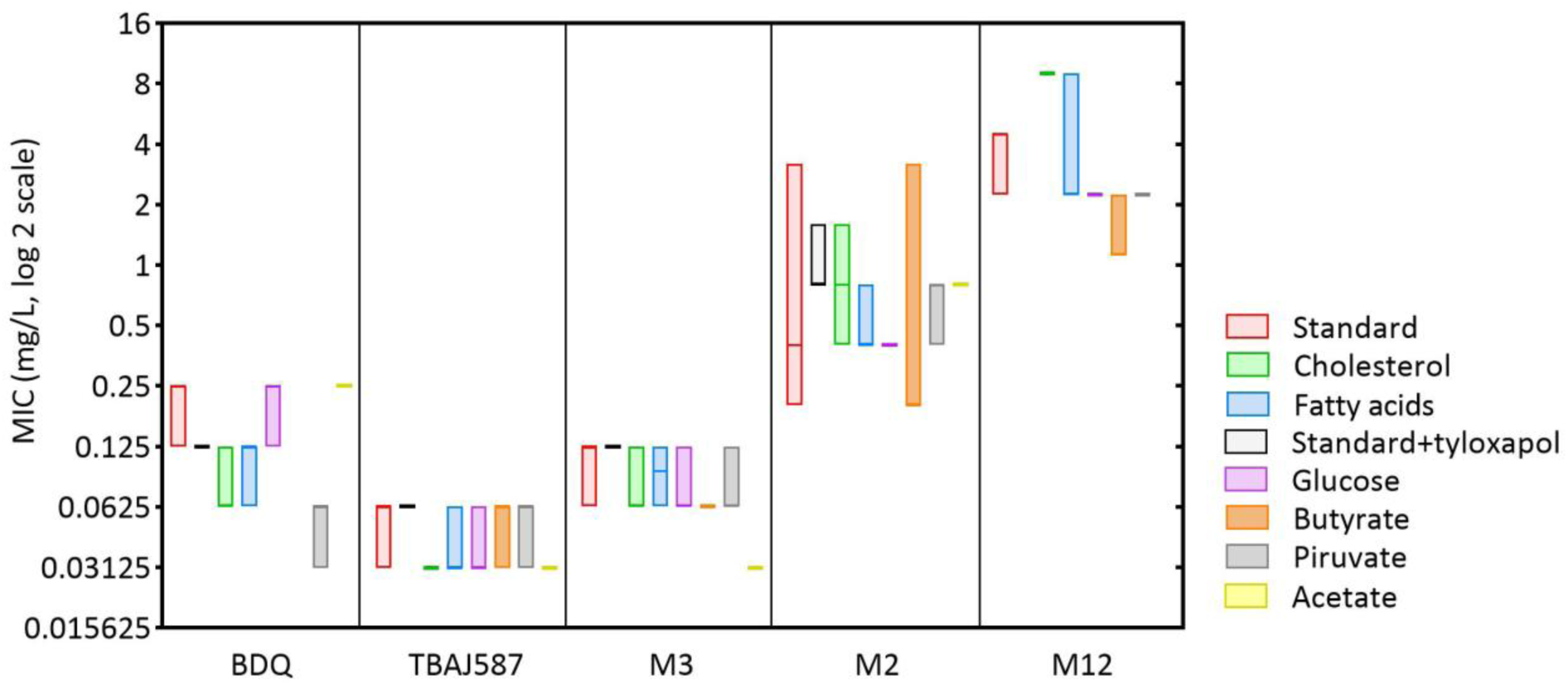
MIC distribution values of TBAJ-587 and its main metabolites against *M. tuberculosis* H37Rv in different broth media. Seven different media compositions with different carbon sources were compared to the standard 7H9 medium, which has dextrose and glycerol as carbon source. MIC values were obtained from at least three technical replicates and plotted in interleaved low-high boxes with line at median. Bedaquiline was included for comparison.

### Longitudinal *in-vitro* pharmacodynamics of TBAJ-587 and its main metabolites against *M. tuberculosis* H37Rv

Longitudinal bacterial survival upon treatment is depicted in **Figure 2**. Untreated growth controls revealed that mycobacterial growth was enhanced in CHO media as compared to FA and ST media. Likewise, for all drug concentrations, higher bacterial loads were observed in CHO treated conditions than in the other two media. In general, when comparing the relative bacterial burden to baseline of the different cultures, the activity of the compounds was higher in ST and FA media compared to CHO medium. Compounds did not have any effect on bacterial growth at concentrations equal and below ½x MIC (MIC_TBAJ587_ = 0.031 mg/L; MIC_M2_= 0.4 mg/L; MIC_M3_= 0.125 mg/L; MIC_M12_= 4.5 mg/L), bacteriostatic effects were observed at 1x MIC and 2x MIC, and bactericidal activities were observed at concentrations equal and above 5x MIC values, suggesting time and concentration dependent activity, i.e., the bactericidal effect of TBAJ-587, M2 and M3 increased progressively from 5x to 300x MIC, especially when treatment was extended over time. Additionally, TBAJ-587 and M3 showed improved activities than M2 and M12 metabolites and bacterial regrowth was observed on day 28 with compounds at 1x MIC and 2x MIC.

**Figure 2.**
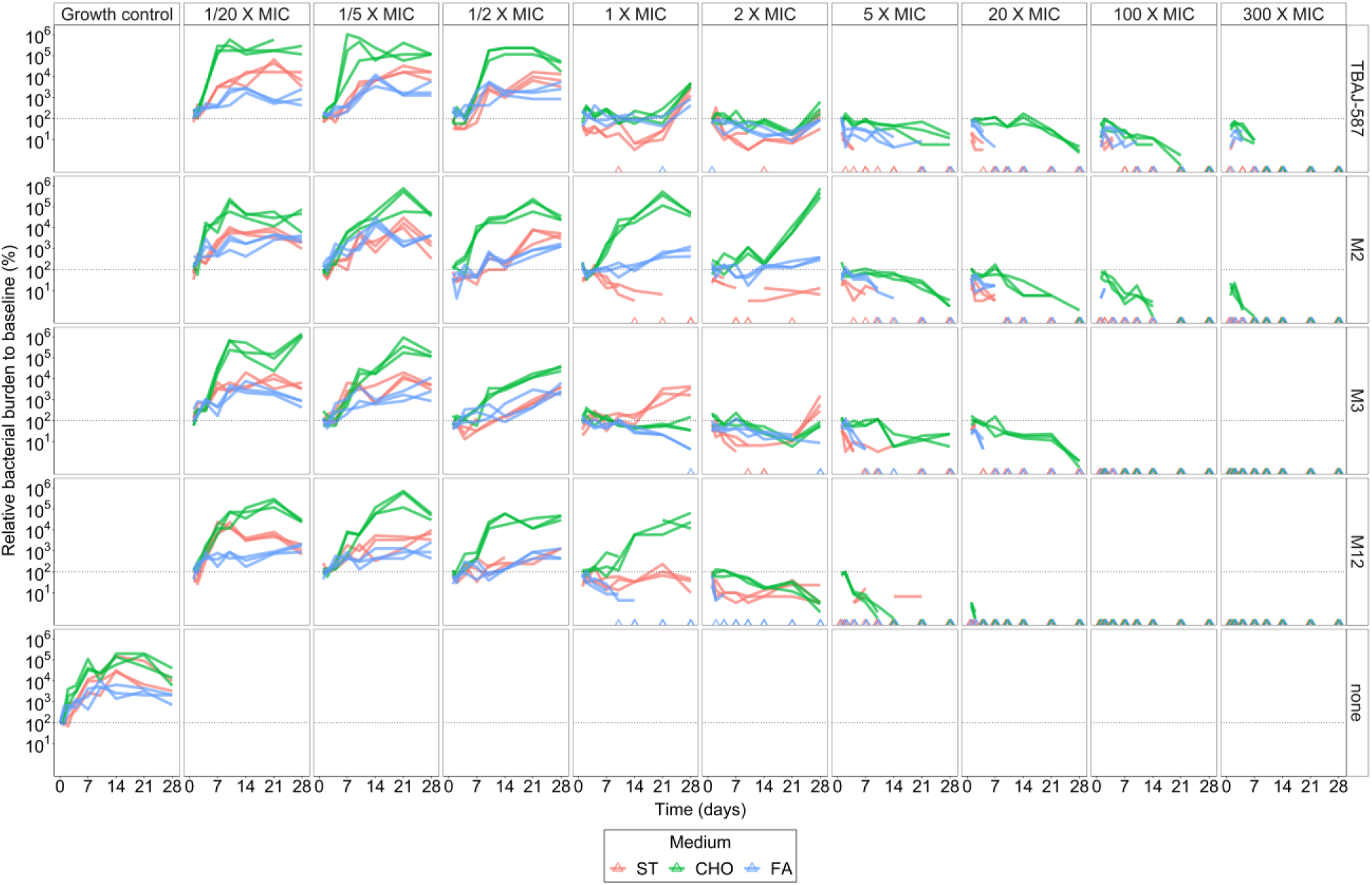
***In vitro* pharmacodynamics of TBAJ-587 and its main metabolites against *M. tuberculosis.*** Bacterial survival of the TKA is stratified by concentration and compound showcasing the relative bacterial burden to baseline (%) of the median CFU/mL in logarithmic scale versus time. Three media were used: standard (ST), cholesterol (CHO) and fatty acids (FA). Triangles indicates counts below quantification limit (bottom of graphs).

Resistance development to TBAJ-587 was investigated in regrowth isolates from ST, CHO, and FA conditions. Following an outgrowth sub-cultivation in ST broth without drug, all isolates exhibited MIC values for TBAJ-587 ranging from 0.125 to 0.25 mg/L (Table S3). This represents a 4-fold increase compared to the wild-type baseline, which suggests emergence of resistance. Whole genome sequencing revealed non-silent mutations across the *Rv0678* gene in the form of deletions, insertions, and single nucleotide polymorphisms (SNP) in all regrowth isolates (**Table 1**). New variants of *Rv0678* mutants were identified in gene codons not reported as associated to resistance before, i.e., R82L and the deletion of AT in the gene position 422-423, truncating codon 141 of the Rv0678 (**Figure 3**). There was a correlation between the highest MIC (0.25 mg/L) in regrowth TKA isolates and the highest frequency of mutations in *Rv0678*. Additionally, two isolates showed low-frequency mutations in *atpB* (L168V) and *atpE* (I66L) genes, while other low-frequency mutations were found in genes not directly associated with drug resistance in *Mtb* (Table S4).

**Table 1.** BDQ-associated resistance mutations in regrowth TKA isolates from this study. MIC values are reported in mg/L.

| Strain ID | xMIC Broth | Genome position | Gene Position | Codon position | Gene | Type | Freq | Mutation | MIC |
| --- | --- | --- | --- | --- | --- | --- | --- | --- | --- |
| 1 | 1X-ST | 779131 | 142 | 47 | *Rv0678* | Deletion | 43.88 | 142 del c* | 0.125 |
| 1 | 1X-ST | 779165 | 176 | 59 | *Rv0678* | Deletion | 14.89 | 176 del c* |  |
| 1 | 1X-ST | 779450 | 461 | 154 | *Rv0678* | SNP | 24.21 | L154P<br>(ctg→cCg) |  |
| 2 | 1X-CHO | 779035-6 | 46-47 | 15 | *Rv0678* | Deletion | 14.81 | 46 del at* | 0.125 |
| 2 | 1X-CHO | 779191 | 202 | 68 | *Rv0678* | SNP | 21.28 | S68G<br>(agc→Ggc) |  |
| 2 | 1X-CHO | 1460745 |  | 168 | *Rv1304*<br>(*atpB*) | SNP | 23.08 | L168V<br>(ctc→Gtc)* |  |
| 3 | 1X-FA | 779363 | 374 | 125 | *Rv0678* | SNP | 5.45 | L125P<br>(ctg→cCg) | 0.125 |
| 4 | 2X-ST | 779411-2 | 422-423 | 141 | *Rv0678* | Deletion | 62.50 | 422 del at* | 0.25 |
| 4 | 2X-ST | 1461240 |  | 166 | *Rv1305*<br>(*atpE*) | SNP | 25.40 | I66L (atc→Ctc)* |  |
| 5 | 2X-CHO | 779192 | 203 | 68 | *Rv0678* | Ins | 17.78 | 203 ins gg* | 0.125 |
| 5 | 2X-CHO | 779216 | 227 | 76 | *Rv0678* | Deletion | 14.74 | 227 del a* |  |
| 6 | 2X-FA | 779234 | 245 | 82 | *Rv0678* | SNP | 88.70 | R82L (cgg/cTg)* | 0.125-0.25 |

\*Asterisks indicate novel mutations identified in this study.

**Figure 3.**
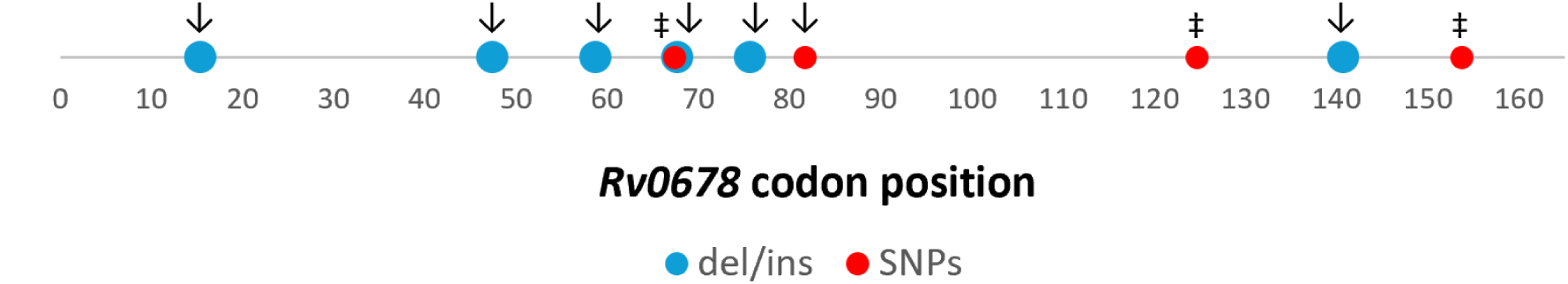
Coding position of mutations identified in *rv0678* found in regrowth isolates from TKA with TBAJ-587. New mutations in *Rv0678* codons associated to DARQ resistance identified in this study (↓) and mutations variants (‡)^35^ previously reported are depicted in Rv0678 codon position.

### Longitudinal *in-vitro* pharmacokinetics of TBAJ-587 and its main metabolites

Measurements of drug concentrations were directly performed in TKA samples with bacteria from six time-points. **Figure 4** represents relative expected (targeted nominal) concentrations to actual measured ones in percentage values. Actual drug quantification over time is displayed in **Figures S1-S12**. Measured experimental concentrations deviated from expected in most culture conditions, which rapidly decreased up to 90% in some cases by day 1 of treatment, at least partially due to unspecific binding to the plastic ware. To test this hypothesis, only M3 metabolite at 10x MIC in standard 7H9 + 0.5% BSA medium was tested by comparing the concentration percentage recovery using glass-tubes or low-binding polypropylene tubes (**Figure S13**). After six days of incubation at 37°C, the recovery in glass tubes was higher than in the polypropylene tubes (83% versus 26%, respectively).

**Figure 4.**
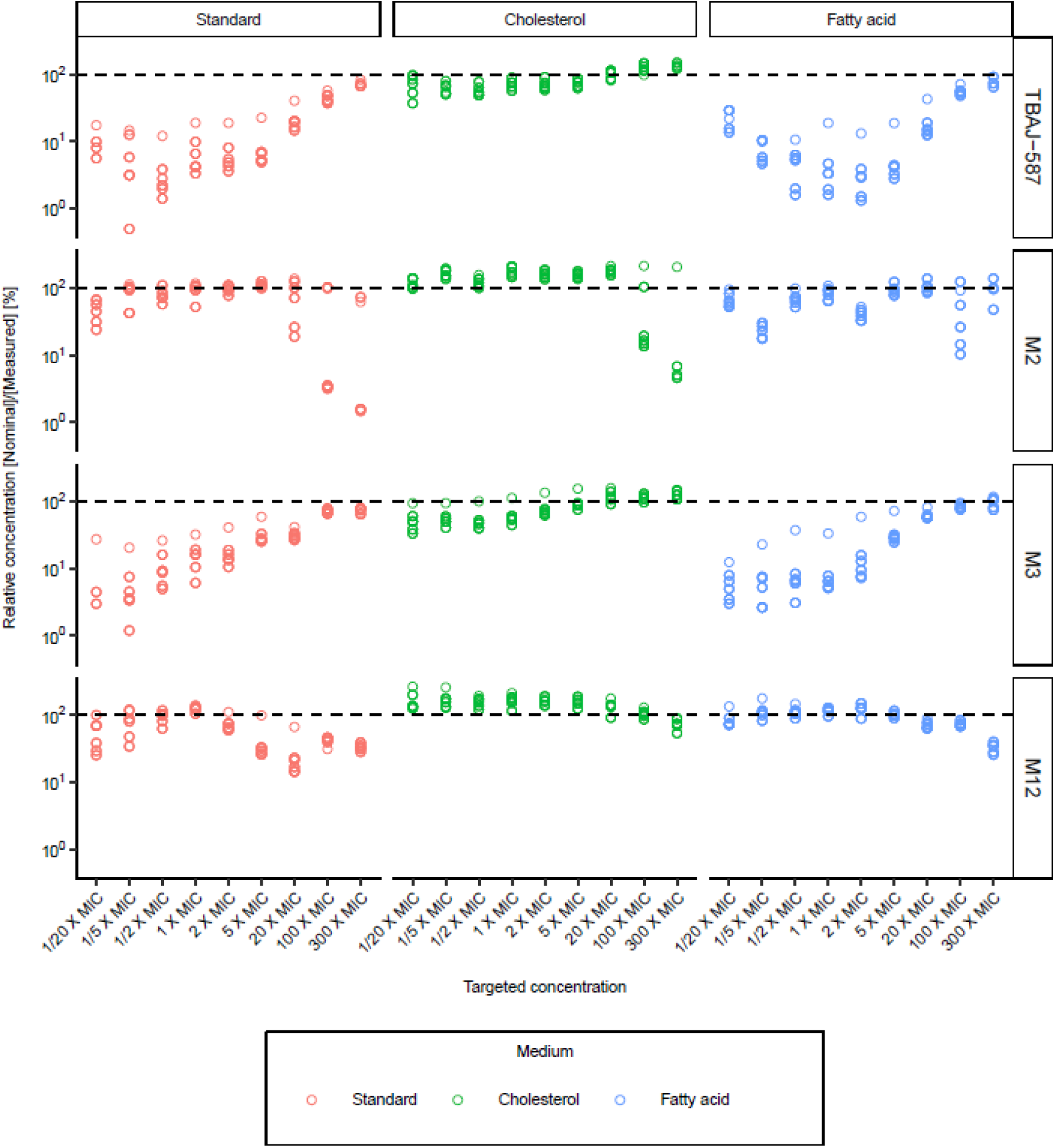
I*n vitro* pharmacokinetics of TBAJ-587 and its main metabolites. Relative expected concentrations (nominal) to actual measured ones in percentage values. Relative concentrations are stratified by concentration and compound. Three media were used: standard, cholesterol and fatty acids. Dashed lines represent the 100% line (relative nominal concentration). Circles represent measured concentrations.

During the TKA, TBAJ-587 and the M3 metabolite exhibited greater deviation from expected concentrations in ST and FA media, particularly at lower concentrations, compared to the M2 and M12 metabolites. After the rapid initial concentration decline observed in day 1, due to plasticware binding, detectable concentrations remained stable throughout the assay, demonstrating that all compounds were stable in the assay media in the presence of bacteria.

A key difference between ST, FA and CHO media is the presence of tyloxapol in the CHO broth, which is included to aid cholesterol solubilization in the assay medium. The influence of this detergent on the stability of the TBAJ-587 compound was investigated in ST broth with and without 0.05% tyloxapol, and in the absence of bacteria. Tyloxapol allowed a higher TBAJ-587 recovery regardless of tested concentrations. Propranolol controls, included in parallel, demonstrated that the presence of tyloxapol did not affect its recovery (**Figure 5**). These data suggest that TBAJ-587 could be solubilized by tyloxapol, thus improving its recovery from the assay media and evidence that nominal concentrations added at the start of the assay do not reflect actual experimental concentrations.

**Figure 5.**
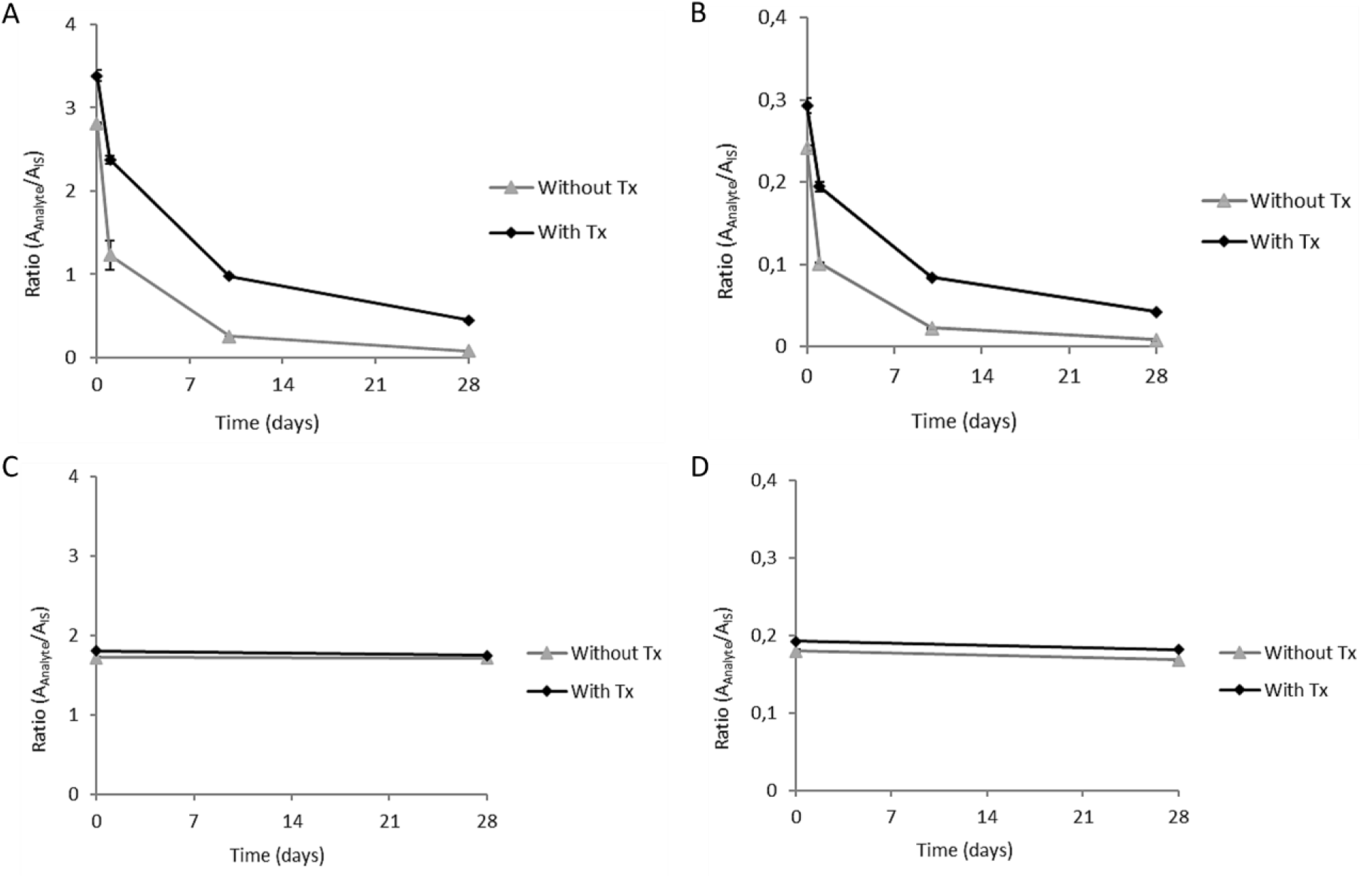
Impact of tyloxapol on the stability of TBAJ-587. Recovery of TBAJ-587 was evaluated over time at **(A)** high (1.6 mg/L) and **(B)** low (0.16 mg/L) concentrations in ST broth medium without and with 0.05 % tyloxapol in the absence of bacteria. The control drug propranolol was also evaluated in the same conditions as TBAJ-587 at **(C)** high (1.6 mg/L) and **(D)** low (0.16 mg/L) concentrations.

### *In vitro* PKPD relationship of TBAJ-587 and its main metabolites

PD (bacterial killing dynamics) and actual *in-vitro* PK (drug stability) measurements in ST, CHO and FA media were integrated by linking the activity of a compound to its effective concentration at every time point. **Figure 6** plots the bacterial burden against both the expected compound exposure (based on the added concentration in flasks) and the actual compound exposure (based on measured concentrations overtime), where better PKPD performances are shown at lower exposure levels, as it is the case of TBAJ-587 in ST and FA cultures, and M3 in FA culture. The mean area under the concentration-time curves (AUC) of actual and expected exposures were then calculated, and ratios (AUC_actual-exposure_/AUC_expected-exposure_) were noted in **Table S5.** Then, the impact of compound binding was assessed on actual PKPD performance, where values close to 1 indicate no meaningful differences in PKPD performance between the actual and the expected drug exposures. Results showed a lower AUC_actual-exposure_ of TBAJ-587 and M3 metabolite in ST and FA media compared with their AUC_expected-exposures_, a pattern not observed or negligible in any other compound-media condition (**Figure 6**). The AUC ratio (actual/expected) of TBAJ-587 was 0.05 and 0.06 for ST and FA, respectively, and 0.15 and 0.07 for M3 metabolite in ST and FA, respectively (**Table S5**); this demonstrates a remarkable difference between expected and actual PKPD performance, suggesting the activity of TBAJ-587 and M3 is underestimated in ST and FA media broths of *in-vitro* assays when PKPD analysis does not account for potential compound binding to plastic ware. In contrast, the same compounds in CHO media broth show AUC ratios of 0.71 and 0.56 for TBAJ-587 and M3, respectively, indicating a higher similarity between the actual and the expected PKPD performance than that seen in ST and FA, although binding was still detected. These findings suggest that bacterial killing was achieved at exposures from 7 to 20-fold times lower of TBAJ-587 (1/0.05 and 1/0.06 for ST and FA, respectively) and M3 (1/0.15 and 1/0.07 for ST and FA, respectively) than theoretically expected, indicating an underestimation of their activities in *in-vitro* assays. ^25^

**Figure 6.**
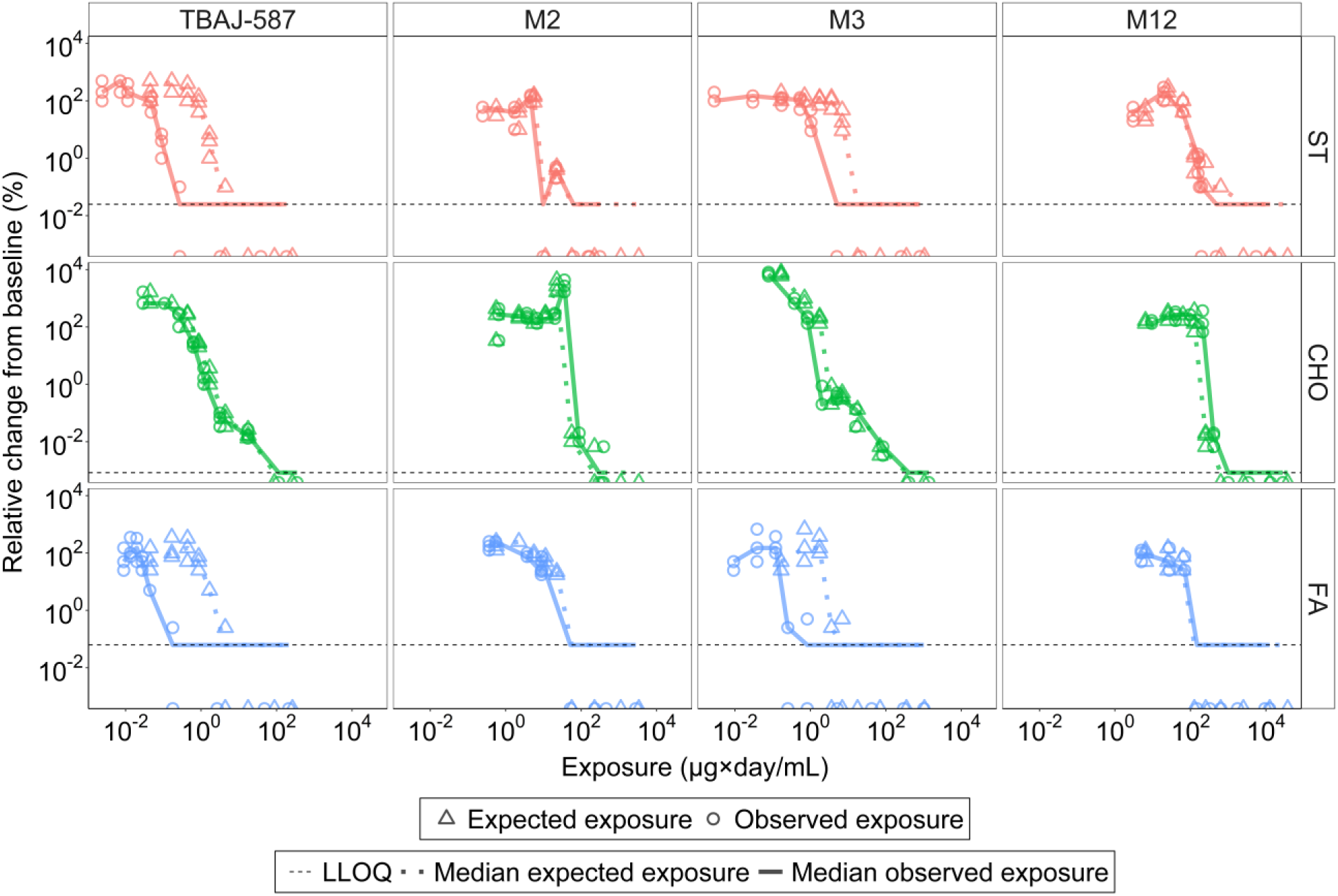
*In vitro* PKPD relationship of TBAJ-compounds in ST, CHO and FA media. Bacterial burdens were plotted against expected and actual compound exposures with curves compared to assess the impact of compound binding. *in-vitro*Bacterial survival, expressed as the percentage of the relative change from baseline versus drug exposure is depicted for expected (dashed lines) and actual observed (solid lines) exposures. ST, standard medium; CHO, cholesterol medium; FA, fatty acid medium. The area under the curve (AUC) of both actual (AUC_actual-exposure_) and expected (AUC_expected-exposure_) was computed and the ratio of actual/expected exposures is noted in table S5.

## DISCUSSION

In 1958, Middlebrook developed synthetic standard broths for drug susceptibility tests (DST) against *Mtb* ^25^, which have been widely used in diagnostics and in preclinical *in-vitro* research. However, there is a need for alternative formulations to better mimic the diverse physiological states and actual drug performance in TB lesions ^26^. Choosing the right carbon source is crucial in drug development, as shown by Pethe K *et al.* ^27^ of the misleading use of glycerol in *in-vitro* media. *Mtb* can utilize fatty acids and cholesterol, which impact its metabolism differently than dextrose and glycerol, the classic carbon sources used in standard 7H9 media ^28^. The use of different carbon sources such as lipids is thus crucial in drug development and it has been shown to impact the activity of compounds and drug combinations^29,30^. This approach targets different bacterial populations, including persisters and dormant forms, which may resist standard treatments.

Our study included eight different broth compositions using alternative carbon sources (ST: dextrose + glycerol; CHO: cholesterol + tyloxapol; FA: oleic, stearic, and palmitic fatty acids; ST + tyl: dextrose + glycerol + tyloxapol; glucose; pyruvate; acetate; and butyrate) in the 7H9-base medium for MIC determinations as a first screening in drug susceptibility. TBAJ-587 demonstrated higher potency as compared to BDQ in six (ST, CHO, FA, ST + tyl, glucose, and acetate) out of the eight media and was the only compound with a minimal MIC shift (2-fold shift across conditions). This indicates that TBAJ-587 had the narrowest range of concentrations to inhibit *Mtb* H37Rv populations developed across all carbon source media, being more potent than BDQ. The M3 metabolite also displayed a better profile than BDQ with a narrower MIC range (**Figure 1**) and, unlike the previously reported activity of the major M2 metabolite of BDQ ^5^, M3 exhibited comparable *in-vitro* activity to TBAJ-587. While M2 and M12 metabolites of TBAJ-587 showed lower activity within a wider range of concentrations across carbon sources than the other DARQs, placing M3 as the most relevant TBAJ-587 metabolite in terms of efficacy against the H37Rv strain in *in-vitro* assays among tested metabolites of this study.

The presence of different lipids in *in-vitro* cultures promote bacterial states different from those found in standard cultures ^30–32^. Our study demonstrated that *Mtb* H37Rv bacterial growth was enhanced in CHO conditions compared to ST and FA, both in treated and untreated conditions. CHO broth includes a non-ionic wetting agent (tyloxapol) to enhance cholesterol solubility, but it also promotes bacterial dispersion, reducing clumping. In contrast, clumping was observed in ST and FA conditions, which could interfere with accurate CFU counts. To address this, collected samples for bacterial measurements were resuspended in PBS with tyloxapol to ensure a homogeneous suspension, which has demonstrated improved CFU enumeration accuracy^33^ to avoid underestimation in ST and FA. Differences in bacterial survival and untreated cultures found in CHO broth in comparison with ST and FA media, suggest that cholesterol broth composition played a role in the increased bacterial burden and survival of the H37Rv strain in the current *in-vitro* TKA with TBAJ-587 and metabolites.

Previous studies demonstrated that the *in-vitro* PKPD driver of BDQ is time-dependent rather than concentration-dependent. *In vitro* TKA using BDQ at 10x and 100x MIC values showed equal efficacy over 12 days of incubation in ST cultures^1^ and this was also observed at 30x MIC and 300x MIC values^34^. However, TBAJ-587 and its M3 metabolite displayed a bacteriostatic effect at concentrations 1x MIC and 2x MIC with subsequent bacterial regrowth by day 28 of the TKA and a bactericidal effect at concentrations above 5x MIC (**Figure 2**). In clinical practice, baseline, treatment initiation, and post-treatment drug susceptibility screening of isolates from patients are essential to detect the potential rapid drug resistance evolution of *Mtb* ^8^. We applied a similar approach in our *in-vitro* TKA in which MICs were determined before and after TKA. We observed a 4-fold shift in the MIC (28 days post-baseline) of cultures treated with 1x and 2x MIC of TBAJ-587, along with different variants associated with resistance across all conditions, regardless of the carbon source (**Table 1**). Known resistance-associated variants of the *Rv0678* gene (L154P, S68G, and L125P found in clinical isolates) were also identified in our study (**Figure 3**), along with new variants in codons linked with resistance (but different type mutation^8,35^). The moderate MIC increase found in isolates from regrowth cultures could be attributable to *Rv0678*, *atpE*, and *atpB* variants. BDQ MIC shifts are documented in *atpE* mutants (from 4-to 128-fold increase), where a moderate shift belongs to strains harboring the variant I66V (4-fold increase)^36^, while a higher fold MIC shift more frequently reported was associated in mutants with the variant I66M ^35,37–39^. In our study, the moderate response to TBAJ-587 in cases with an *atpE* mutation may be attributable to the specific amino acid substitution (I66L, here first described). It is not clear if mutations in other genes found in regrowth cultures (*mce3A*, *rfbD*, *hsdM*, and *mgtA*, *ercc3*, *uvrB*, *sigM*, *argF*, and *cyp137*, **Table S4**) could be related to *Mtb* pharmacodynamics response (MIC and TKA) to TBAJ-587, as they are more often associated with survival mechanisms like DNA repair, oxidative stress, cell wall transportation/biosynthesis, etc., rather than resistance. Thus, the use of different carbon sources in *in-vitro* drug susceptibility assays allowed the identification of a wider range of populations associated with phenotypic resistance.

For some compounds, such as beta-lactams, bacterial regrowth in *in-vitro* time-kill assays is typically associated with drug instability in the assay media, which can be misleading when interpreting true antibacterial activity^40^. In the case of TBAJ-587 and its main metabolites, bacterial regrowth at 1x MIC and 2x MIC was not associated to drug instability (**Figure 4**). Although measured experimental concentrations were lower than theoretically expected ones, markedly in ST and FA cultures, concentrations sharply decreased at the first time-point of quantification; after which, these remained stable for the rest of the time course under all culture conditions. This pattern would be uncommon if the compound were unstable due to degradation, whith a gradually decline of drug concentration with a time to half-life. We thus hypothesize that, due to their lipophilic nature, TBAJ--587 and its metabolites exhibit adsorption to the flask material, polystyrene, resulting in an unbound fraction that remained stable over the 28 days of the TKA. Yet, remaining drug concentrations in solution were effective enough to kill bacteria overtime, suggesting that *in-vitro* potency of the TBAJ-587 compounds may be underestimated in assays where compound stability is not confirmed. Likewise, variations in drug concentrations observed as time progressed can be attributed to reversible binding interactions between TBAJ-587 compound and the plastic material.

Non-specific adsorption of drugs to plastics is a phenomenon influenced by the physicochemical properties of compounds, the solvent composition, the time of contact and temperature, which reduces the drug concentration in solution ^41^. Previous studies suggest that BDQ has stronger interactions with polypropylene than with polystyrene lab-wares attributed to its high lipophilicity. Lounis *et al.* ^20^ showed a 4-fold increase in the MIC of BDQ when using polypropylene plates in comparison with polystyrene, thus confirming the plastic binding hypothesis. Our observations clearly indicate that TBAJ-compounds bind polystyrene lab-wares, though with a likely lower affinity than polypropylene materials. The adsorption effect was mainly observed when using ST and FA broth and negligible in CHO medium. CHO broth contains tyloxapol, which improved the solubility of TBAJ-587 compounds in solution as demonstrated in drug measurement assays with and without tyloxapol (**Figure 5**), suggesting that the surfactant may reduce the adsorption of compounds to the plastic ware. Although higher drug concentrations in CHO cultures due to less binding than that observed in ST and FA, this did not translate into improved potency of TBAJ-587 and metabolites (**Figure 6**), which could be attributed to the specific metabolism *Mtb* bacilli developed in CHO cultures that may deserve further investigation by transcriptomic studies.

Modest MIC increments of DARQ are associated with mutations in *Rv0678* ^16^, a mutation that confers cross resistance between DARQs and clofazimine ^42^. Identification of concentrations that prevent emergence of resistance is crucial to adjust therapeutic doses for successful clinical outcomes. Unlike BDQ, TBAJ-587 seems to better overcome issues related to modest MIC shifts in the clinic due to its greater potency and safer properties and capacity to prevent emergence of resistance ^16^. Previous *in-vitro* studies reported emergence of resistance within 3 – 6 weeks upon BDQ treatment with 4x MIC and 8x MIC^1^; here we show that treatment with concentrations equal or higher than 5x MIC of TBAJ-587 cleared the cultures by day 7 (bacterial counts under the limit of quantification) without bacterial regrowth after 28 days of treatment. These data mimic mouse model observations, to some extent, in which TBAJ-587 exhibited reduced selection of resistance, compared to BDQ, when administrated in monotherapy and in combination with pretomanid with linezolid (PaL) or pretomanid with moxifloxacin and pyrazinamide (PaMZ)^16^.

In summary, in this study we integrated *in-vitro* longitudinal studies with actual PK and PD parameters to provide actual exposures of TBAJ-587 and its metabolites against the reference *Mtb* H37Rv strain in different carbon sources, showing that their actual *in-vitro* activity might have been underestimated, since assays regularly reported do not consider drug stability. This observation will feed translational *in silico* PKPD models to refine dose optimization. Our *in-vitro* study further supports the progression of the TBAJ-587 in the drug development pipeline for TB, establishing initial *in-vitro* PKPD cut-offs for prevention of emergence of resistance.

## ACKNOWLEDGEMENTS

Authors would like to acknowledge the use of Servicio General de Apoyo a la Investigación-SAI, Universidad de Zaragoza.

We would like to thank Begoña Gracia Díaz and Aleksandra Muzylo from the University of Zaragoza for their technical and lab management support.

This work has received support from the Innovative Medicines Initiatives 2 Joint Undertaking (grant No 853989).

## TRANSPARENCY DECLARATIONS

Natalya Serbina is the Senior Director of Biology at TB Alliance. The rest of the authors have none to declare.

## AUTHOR CONTRIBUTION

CrediT (Contributor Roles Taxonomy) has been applied for author contribution. Conceptualization: NS, AL, SRG; Data curation: DAAA, MREM, AAML; Formal Analysis: DAAA, AAML. LS; Funding acquisition: NW, USHS, SRG; Investigation: DAAA, MSR, MREM, APM; Methodology: DAAA, MREM, AAML, SRG; Project administration: NS, USHS, AL, SRG; Resources: NW, NS, USHS, LS, AAML, SRG; Supervision: NW, NS, USHS, AL, SRG; Visualization: DAAA, MREM, AAML; Writing – original draft: DAAA, SRG; Writing – review & editing: DAAA, AAML, NS, NW, LS, MREM, NW, MSR, SRG. All authors read and approved the final version of the document.

