## Supplementary Tables S1-S5, and Figures S1-S13 for "*In vitro* pharmacokinetics and pharmacodynamics of the diarylquinoline TBAJ-587 and its metabolites against *Mycobacterium tuberculosis*"

**Running title (52/54 characters):** *In vitro* PKPD of TBAJ-587 for Tuberculosis treatment

**Keywords:** tuberculosis, TBAJ-587, pharmacokinetics/pharmacodynamics, time-kill assay.

### SUPPLEMENTARY TABLES

**Table S1. Time-kill assays conditions for TBAJ-587 and its main metabolites against *M. tuberculosis* H37Rv.**

| Condition Num. | Drug | Media | Concentration (mg/L) | Specifications |
|---|---|---|---|---|
| 1/2/3 | TBAJ587 | ST/CHO/FA | 0,0016 | Ca. 1/20x MIC |
| 4/5/6 | TBAJ587 | ST/CHO/FA | 0,006 | Ca. 1/5x MIC |
| 7/8/9 | TBAJ587 | ST/CHO/FA | 0,016 | Ca. 1/2x MIC |
| 10/11/12 | TBAJ587 | ST/CHO/FA | 0,031 | Ca. 1x MIC |
| 13/14/15 | TBAJ587 | ST/CHO/FA | 0,063 | Ca. 2x MIC |
| 16/17/18 | TBAJ587 | ST/CHO/FA | 0,156 | Ca. 5x MIC |
| 19/20/21 | TBAJ587 | ST/CHO/FA | 0,625 | Ca. 20x MIC |
| 22/23/24 | TBAJ587 | ST/CHO/FA | 3,13 | Ca. 100x MIC |
| 25/26/27 | TBAJ587 | ST/CHO/FA | 9,38 | Ca. 300x MIC |
| 28/29/30 | M2 | ST/CHO/FA | 0,02 | Ca. 1/20x MIC |
| 31/32/33 | M2 | ST/CHO/FA | 0,08 | Ca. 1/5x MIC |
| 34/35/36 | M2 | ST/CHO/FA | 0,2 | Ca. 1/2x MIC |
| 37/38/39 | M2 | ST/CHO/FA | 0,4 | Ca. 1x MIC |
| 40/41/42 | M2 | ST/CHO/FA | 0,8 | Ca. 2x MIC |
| 43/44/45 | M2 | ST/CHO/FA | 2 | Ca. 5x MIC |
| 46/47/48 | M2 | ST/CHO/FA | 8 | Ca. 20x MIC |
| 49/50/51 | M2 | ST/CHO/FA | 40 | Ca. 100x MIC |
| 52/53/54 | M2 | ST/CHO/FA | 120 | Ca. 300x MIC |
| 55/56/57 | M3 | ST/CHO/FA | 0,006 | Ca. 1/20x MIC |
| 58/59/60 | M3 | ST/CHO/FA | 0,025 | Ca. 1/5x MIC |
| 61/62/63 | M3 | ST/CHO/FA | 0,063 | Ca. 1/2x MIC |
| 64/65/66 | M3 | ST/CHO/FA | 0,13 | Ca. 1x MIC |
| 67/68/69 | M3 | ST/CHO/FA | 0,25 | Ca. 2x MIC |
| 70/71/72 | M3 | ST/CHO/FA | 0,63 | Ca. 5x MIC |
| 73/74/75 | M3 | ST/CHO/FA | 2,5 | Ca. 20x MIC |
| 76/77/78 | M3 | ST/CHO/FA | 12,5 | Ca. 100x MIC |
| 79/80/81 | M3 | ST/CHO/FA | 37,5 | Ca. 300x MIC |
| 82/83/84 | M12 | ST/CHO/FA | 0,23 | Ca. 1/20x MIC |
| 85/86/87 | M12 | ST/CHO/FA | 0,9 | Ca. 1/5x MIC |
| 88/89/90 | M12 | ST/CHO/FA | 2,25 | Ca. 1/2x MIC |
| 91/92/93 | M12 | ST/CHO/FA | 4,5 | Ca. 1x MIC |
| 94/95/96 | M12 | ST/CHO/FA | 9 | Ca. 2x MIC |
| 97/98/99 | M12 | ST/CHO/FA | 22,5 | Ca. 5x MIC |
| 100/101/102 | M12 | ST/CHO/FA | 90 | Ca. 20x MIC |
| 103/104/105 | M12 | ST/CHO/FA | 450 | Ca. 100x MIC |
| 106/107/108 | M12 | ST/CHO/FA | 1350 | Ca. 300x MIC |
| 109/110/111 | None | ST/CHO/FA | No drug | Growth controls |

**Table S2. MIC determination of TBAJ-587 and its main metabolites against *M. tuberculosis* H37Rv Pasteur in eight different carbon sources.** At least three technical replicates were carried out for each determination

| Compound/ Medium | MIC (mg/L) | | | | | | | |
|---|---|---|---|---|---|---|---|---|
| | **Standard** | **Cholesterol + tyloxapol** | **Standard + tyloxapol** | **Fatty acids** | **Glucose** | **Pyruvate** | **Acetate** | **Butyrate** |
| TBAJ587 | 0.062<br>0.062<br>0.031<br>0.062<br>0.062<br>0.031<br>0.062<br>0.031<br>0.031 | 0.031<br>0.031<br>0.031<br>0.031<br>0.031<br>0.031 | 0.062<br>0.062<br>0.062 | 0.062<br>0.062<br>0.031<br>0.031<br>0.031<br>0.031 | 0.031<br>0.031<br>0.062 | 0.031<br>0.062<br>0.062 | 0.031<br>0.031<br>0.031 | 0.062<br>0.062<br>0.031 |
| M2 | 0.4<br>0.4<br>0.2<br>0.4<br>0.4<br>3.2<br>0.4<br>0.4<br>0.4 | 0.8<br>0.8<br>0.4<br>0.8<br>1.6<br>1.6 | 0.8<br>0.8<br>1.6 | 0.8<br>0.4<br>0.4<br>0.4<br>0.4<br>0.4 | 0.4<br>0.4<br>0.4 | 0.4<br>0.8<br>0.8 | 0.8<br>0.8<br>0.8 | 0.2<br>0.2<br>3.2 |
| M3 | 0.125<br>0.062<br>0.062<br>0.125<br>0.125<br>0.125<br>0.125<br>0.125<br>0.062 | 0.125<br>0.062<br>0.062<br>0.125<br>0.062<br>0.062 | 0.125<br>0.125<br>0.125 | 0.062<br>0.125<br>0.125<br>0.062<br>0.062<br>0.125 | 0.062<br>0.062<br>0.125 | 0.062<br>0.062<br>0.125 | 0.031<br>0.031<br>0.031 | 0.062<br>0.062<br>0.062 |
| M12 | >2.25<br>>2.25<br>>2.25<br>4.5<br>2.25<br>2.25<br>4.5<br>4.5<br>4.5 | >2.25<br>>2.25<br>>2.25<br>9<br>9<br>9 | >2.25<br>>2.25<br>>2.25 | >2.25<br>>2.25<br>2.25<br>9<br>2.2<br>2.25 | 2.25<br>2.25<br>>2,25 | 2.25<br>2.25<br>2.25 | >2.25<br>>2.25<br>>2.25 | 1.125<br>1.125<br>2.25 |
| BDQ | 0.125<br>0.25<br>0.25 | 0.062<br>0.062<br>0,125 | 0.125<br>0.125<br>0.125 | 0.062<br>0.125<br>0.125 | 0.125<br>0.25<br>0.25 | 0.031<br>0.062<br>0.062 | 0.25<br>0.25<br>0.25 | |
| MXF | 0.125<br>0.125<br>0.25 | 0.062<br>0.062<br>0.062 | 0.062<br>0.062<br>0.125 | 0.062<br>0.062<br>0.125 | 0.062<br>0.125<br>0.25 | 0.125<br>0.125<br>0.125 | 0.125<br>0.125<br>0.125 | |
| LZD | 0.5<br>1<br>2 | 2<br>2<br>2 | 2.0<br>2.0<br>2.0 | 1<br>1<br>2 | 1<br>1<br>2 | 1<br>2<br>2 | 2<br>2<br>2 | |

MIC determinations were performed in triplicate; biological replicates are separated by gray lines. “>”, MIC above the maximum concentration tested. Bedaquiline (BDQ) was included for comparison, and moxifloxacin (MXF) and linezolid (LZD) as internal controls. The concentration range tested for TBAJ-587, M3 and BDQ was 0.002 - 0.25 mg/L. The range for M2 metabolite was 0.012 – 1.6 mg/L. Two ranges were tested for M12: 0.017 – 2.25 mg/L and 0.017 - 18 mg/L.

**Table S3. TBAJ-587 MIC of bacterial regrowth isolates from conditions treated with 1x MIC and 2x MIC values in TKA.** Condition: it is labeled with the condition number, medium broth, the xMIC drug concentration in TKA, and days of incubation in TKA when isolated; MIC: Minimum Inhibitory Concentration carried out in triplicate by REMA in ST broth.

| Isolate | Condition | MIC |
|---|---|---|
| 1 | 10_ST / 1X TBAJ-587 / 28 days | 0.125 |
| 2 | 11_CHO / 1X TBAJ-587 / 28 days | 0.125 |
| 3 | 12_FA / 1X TBAJ-587 / 28 days | 0.125 |
| 4 | 13_ST / 2X TBAJ-587 / 28 days | 0.250 |
| 5 | 14_CHO / 2X TBAJ-587 / 28 days | 0.125 |
| 6 | 15_FA / 2X TBAJ-587 / 28 days | 0.125 - 0.250 |

**Table S4. Whole Genome Sequencing data of the TKA regrowth isolates from this study.**

| Strain ID | xMIC Broth | Position | Ref. | Alternative Allele | Relevance | Category | Gene | Gene name | Type | Cov-For | Cov-Rev | Qual 20 | Freq | Cov | Subst |
|---|---|---|---|---|---|---|---|---|---|---|---|---|---|---|---|
| 1 | 1X-ST | 779131 | C | GAP | resistant | nonessential | *Rv0678* | - | Deletion | 23 | 20 | 43 | 43.88 | 98 | 142 del c |
| 1 | 1X-ST | 779165 | C | GAP | resistant | nonessential | *Rv0678* | - | Deletion | 9 | 5 | 14 | 14.89 | 94 | 176 del c |
| 1 | 1X-ST | 779450 | T | C | resistant | nonessential | *Rv0678* | - | SNP | 15 | 8 | 23 | 24.21 | 95 | L154P (ctg/cCg) |
| 1 | 1X-ST | 1474955 | T | C | unknown | - | *Rvnr02* | *rrl* | SNP | 3 | 9 | 12 | 27.27 | 44 | - |
| 1 | 1X-ST | 2210336 | T | G | unknown | nonessential | *Rv1966* | *mce3A* | SNP | 2 | 7 | 9 | 11.84 | 76 | L337R (ctg/cGg) |
| 1 | 1X-ST | 2210339 | C | A | unknown | nonessential | *Rv1966* | *mce3A* | SNP | 2 | 7 | 9 | 12.16 | 74 | P338Q (ccg/cAg) |
| 1 | 1X-ST | 4229807 | A | G | unknown | nonessential | *Rv3783* | *rfbD* | SNP | 6 | 6 | 12 | 12.00 | 100 | T184A (acc/Gcc) |
| 1 | 1X-ST | 4229813 | T | C | unknown | nonessential | *Rv3783* | *rfbD* | SNP | 6 | 6 | 12 | 11.54 | 104 | Y186H (tac/Cac) |
| 2 | 1X-CHO | 779035-6 | AT | GAP | resistant | nonessential | *Rv0678* | - | Deletion | 7 | 5 | 16 | 14.81 | 81 | 46 del at |
| 2 | 1X-CHO | 779191 | A | G | resistant | nonessential | *Rv0678* | - | SNP | 10 | 10 | 20 | 21.28 | 94 | S68G (agc/Ggc) |
| 2 | 1X-CHO | 1460745 | C | G | resistant | essential | *Rv1304* | *atpB* | SNP | 10 | 8 | 18 | 23.08 | 78 | L168V (ctc/Gtc) |
| 2 | 1X-CHO | 2210336 | T | G | unknown | nonessential | *Rv1966* | *mce3A* | SNP | 3 | 11 | 14 | 17.28 | 81 | L337R (ctg/cGg) |
| 2 | 1X-CHO | 2210339 | C | A | unknown | nonessential | *Rv1966* | *mce3A* | SNP | 3 | 11 | 14 | 17.95 | 78 | P338Q (ccg/cAg) |
| 3 | 1X-FA | 234475 | T | G | unknown | nonessential | *Rv0197* | - | SNP | 4 | 2 | 6 | 20.00 | 30 | Y749D (tat/Gat) |
| 3 | 1X-FA | 234476 | A | C | unknown | nonessential | *Rv0197* | - | SNP | 4 | 2 | 6 | 15.79 | 38 | Y749S (tat/tCt) |
| 3 | 1X-FA | 234478 | C | A | unknown | nonessential | *Rv0197* | - | SNP | 4 | 2 | 6 | 15.00 | 40 | P750T (ccc/Acc) |
| 3 | 1X-FA | 779363 | T | C | resistant | nonessential | Rv0678 | - | SNP | 1 | 2 | 3 | 5.45* | 55 | L125P (ctg/cCg) |
| 3 | 1X-FA | 3068836 | C | T | unknown | nonessential | *Rv2756c* | *hsdM* | SNP | 4 | 4 | 8 | 12.90 | 62 | R416R (cgg/cgA) |
| 4 | 2X-ST | 234475 | T | G | unknown | nonessential | *Rv0197* | - | SNP | 3 | 4 | 7 | 19.44 | 36 | Y749D (tat/Gat) |
| 4 | 2X-ST | 234476 | A | C | unknown | nonessential | *Rv0197* | - | SNP | 3 | 4 | 7 | 16.67 | 42 | Y749S (tat/tCt) |
| 4 | 2X-ST | 234478 | C | A | unknown | nonessential | *Rv0197* | - | SNP | 4 | 4 | 8 | 22.22 | 36 | P750T (ccc/Acc) |
| 4 | 2X-ST | 649106 | T | A | unknown | essential | *Rv0557* | *mgtA* | SNP | 2 | 4 | 6 | 10.00 | 60 | S191T (tcg/Acg) |
| 4 | 2X-ST | 779411-2 | AT | GAP | resistant | nonessential | *Rv0678* | - | Deletion | 24 | 26 | 50 | 62.50 | 80 | 422 del at |

| | | | | | | | | | | | | | | | |
|---|---|---|---|---|---|---|---|---|---|---|---|---|---|---|---|
| 4 | 2X-ST | 906993 | G | A | unknown | nonessential | *Rv0812* | - | SNP | 4 | 4 | 8 | 10.13 | 79 | A191T (gcc/Acc) |
| 4 | 2X-ST | 958563 | A | G | unknown | nonessential | *Rv0861c* | *ercc3* | SNP | 3 | 3 | 6 | 13.33 | 45 | I530T (atc/aCc) |
| 4 | 2X-ST | 990399 | A | T | unknown | nonessential | *Rv0890c* | - | SNP | 3 | 4 | 7 | 14.00 | 50 | L733Q (ctg/cAg) |
| 4 | 2X-ST | 1461240 | A | C | resistant | essential | *Rv1305* | *atpE* | SNP | 10 | 6 | 16 | 25.40 | 63 | I66L (atc/Ctc) |
| 4 | 2X-ST | 1837303 | A | C | unknown | nonessential | *Rv1633* | *uvrB* | SNP | 3 | 3 | 6 | 11.54 | 52 | N77H (aac/Cac) |
| 4 | 2X-ST | 4400580 | A | G | unknown | nonessential | *Rv3911* | *sigM* | SNP | 2 | 2 | 4 | 10.26 | 39 | Q132R (cag/cGg) |
| 5 | 2X-CHO | 779192 | G | C | resistant | nonessential | *Rv0678* | - | Ins | 9 | 7 | 16 | 17.78 | 16 | 203 ins gg |
| 5 | 2X-CHO | 779216 | A | GAP | resistant | nonessential | *Rv0678* | - | Deletion | 9 | 5 | 15 | 14.74 | 95 | 227 del a |
| 5 | 2X-CHO | 1870075 | C | A | unknown | essential | *Rv1656* | *argF* | SNP | 5 | 5 | 10 | 11.49 | 87 | R52S (cgc/Agc) |
| 5 | 2X-CHO | 2181393 | T | A | unknown | nonessential | *Rv1929c* | - | SNP | 2 | 2 | 4 | 16.67 | 24 | T172S (acg/Tcg) |
| 5 | 2X-CHO | 3391069 | G | A | unknown | essential | *Rv3031* | - | SNP | 6 | 2 | 8 | 12.12 | 66 | G383E (gga/gAa) |
| 5 | 2X-CHO | 3391072 | T | C | unknown | essential | *Rv3031* | - | SNP | 6 | 2 | 8 | 11.59 | 69 | F384S (ttc/tCc) |
| 5 | 2X-CHO | 4128130 | T | A | unknown | nonessential | *Rv3685c* | *cyp137* | SNP | 7 | 2 | 9 | 10.84 | 83 | D199V (gac/gTc) |
| 6 | 2X-FA | 779234 | G | T | resistant | nonessential | *Rv0678* | - | SNP | 58 | 44 | 102 | 88.70 | 115 | R82L (cgg/cTg) |

**Criteria for selection of reads with mutations:** reads are non-repetitive genes only, two reads in both strands forward and reverse direction, mutation frequency equal of above 10% (*except for isolate 3, for which not relevant genes were found at higher than 10%), location in a coding gene, and non-synonymous SNP. **Column content: Strain ID** stands for the number of isolate; **Position** indicates the reference nucleotide in genome; **Ref**. indicates the reference allele in the wild type *Mycobacterium tuberculosis* strain; **Alternative Allele** refers to the mutated allele in strains; **Relevance** indicates the relevance of the mutation in the context of resistance to bedaquiline; **Category** refers to the gene's role; **Gene** and **Gene name** are the specific identifier and name, respectively, in the genome of *M. tuberculosis*; **Type** indicates the genetic variation, SNP or Deletion; **Cov-For** refers to the coverage of the forward strand in sequencing; **Cov-Rev** refers to the coverage of the reverse strand in sequencing; **Qual20** is the quality score; **Freq** represents the frequency of the mutation observed; **Cov** refers to the overall coverage; **Subst** describes the specific type of mutation (insertion, deletion or substitution) at given position. Yellow cells highlight variants associated with resistance.

**Table S5. Ratios of the actual exposure relative to expected exposure of TBAJ-587 and its main metabolites.** Exposure based on the area under the concentration-time curve from either measured concentration over time (actual exposure) or the expected concentrations over time (added concentration in culture flasks, expected exposure). AUC-ratios where values close to 1 indicate no meaningful differences in PKPD performance between the actual and the expected drug exposures.

$Relative\ exposure\ = \frac{AUC\_Actual\_exposure}{AUC\_Expected\_exposure}$ ST: standard broth culture, CHO: cholesterol broth culture, FA: oleic, palmitic, and stearic fatty acids broth culture.

| **Compound** | **ST** | **CHO** | **FA** |
|---|---|---|---|
| **TBAJ-587** | 0.05 | 0.71 | 0.06 |
| **M2** | 0.80 | 1.59 | 0.65 |
| **M3** | 0.15 | 0.56 | 0.07 |
| **M12** | 0.69 | 1.58 | 1.05 |

### SUPPLEMENTARY FIGURES

**Figure S1. Drug concentration of TBAJ-587 in standard broth.** Relative concentration to time zero of TBAJ-587 in standard medium for the time kill assay, dotted line marks the baseline concentration. Below quantification limit marked on x-axis.

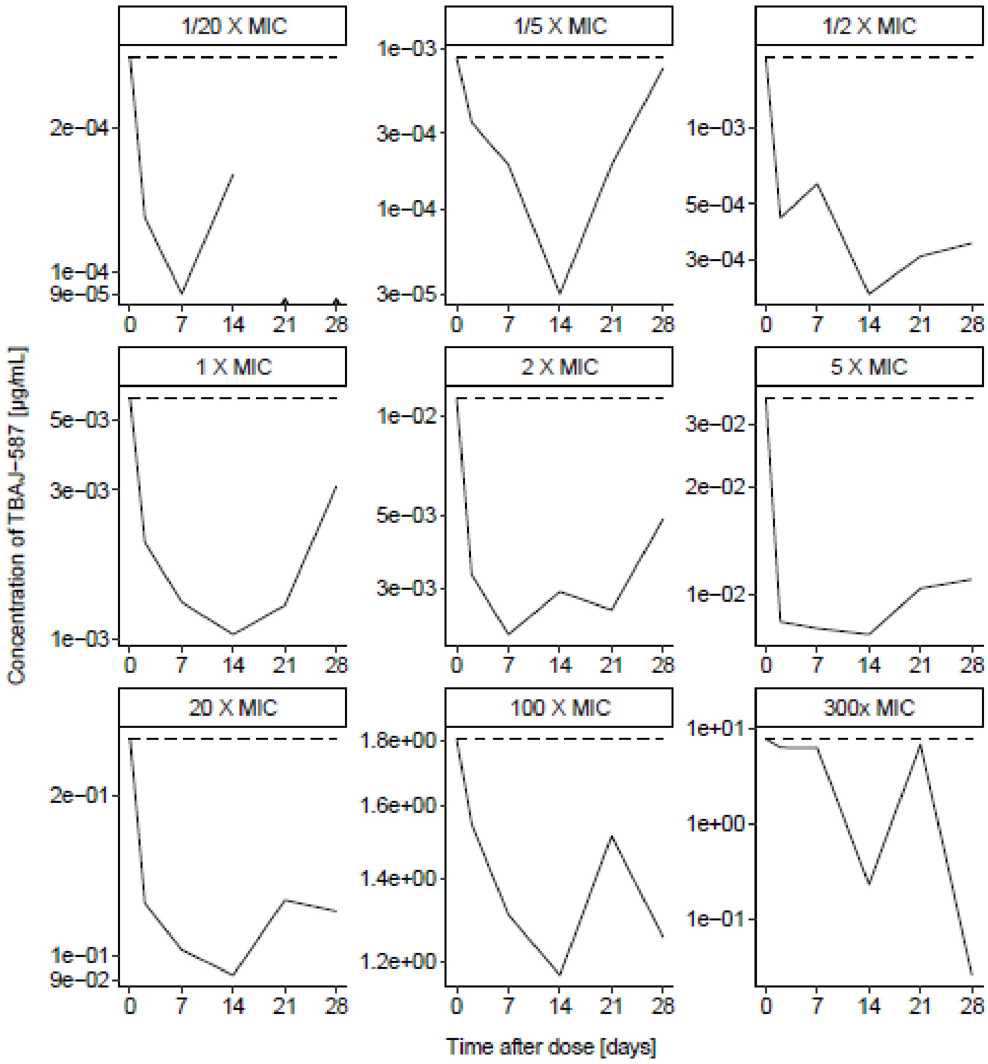

**Figure S2. Drug concentration of TBAJ-587 in cholesterol broth.** Relative concentration to time zero of TBAJ-587 in cholesterol medium for the time kill assay, dotted line marks the baseline concentration. Below quantification limit marked on x-axis.

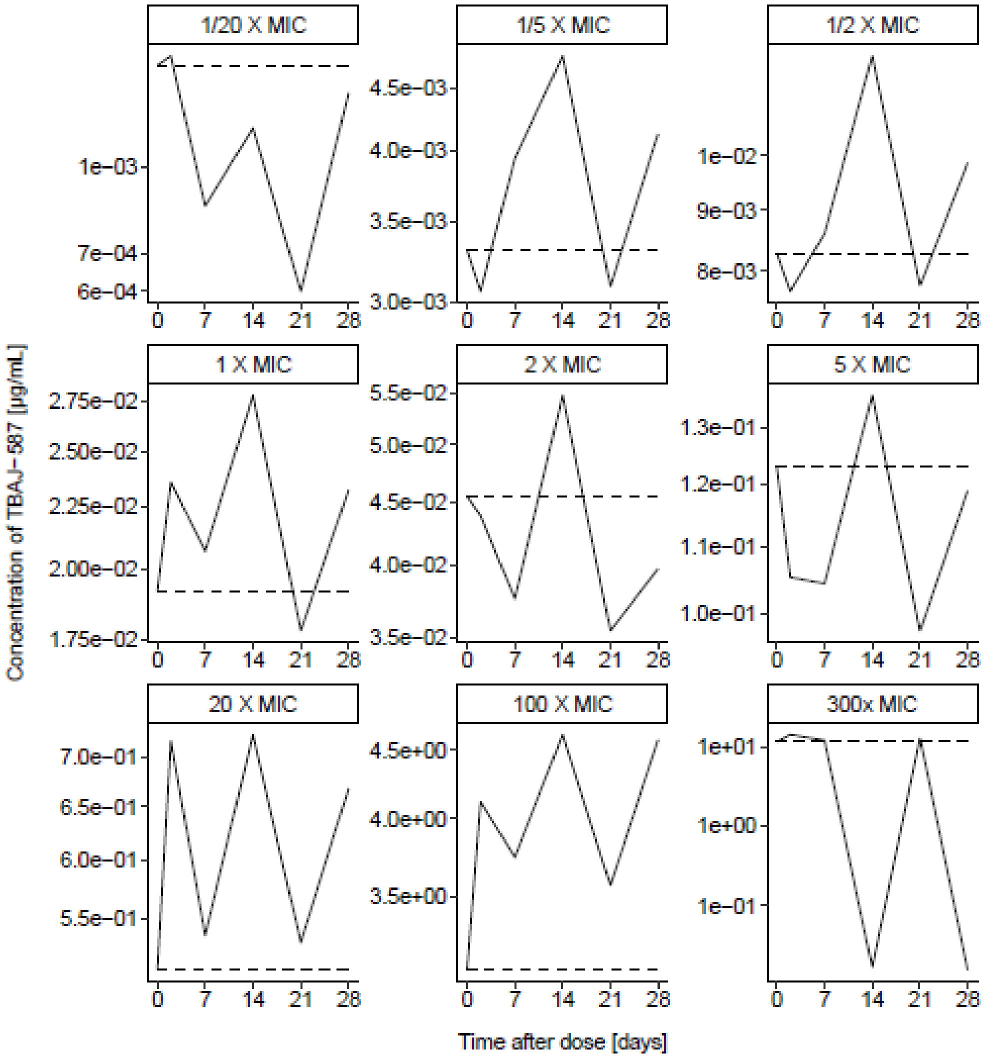

**Figure S3. Drug concentration of TBAJ-587 in fatty acids broth.** Relative concentration to time zero of TBAJ-587 in fatty acids medium for the time kill assay, dotted line marks the baseline concentration. Below quantification limit marked on x-axis.

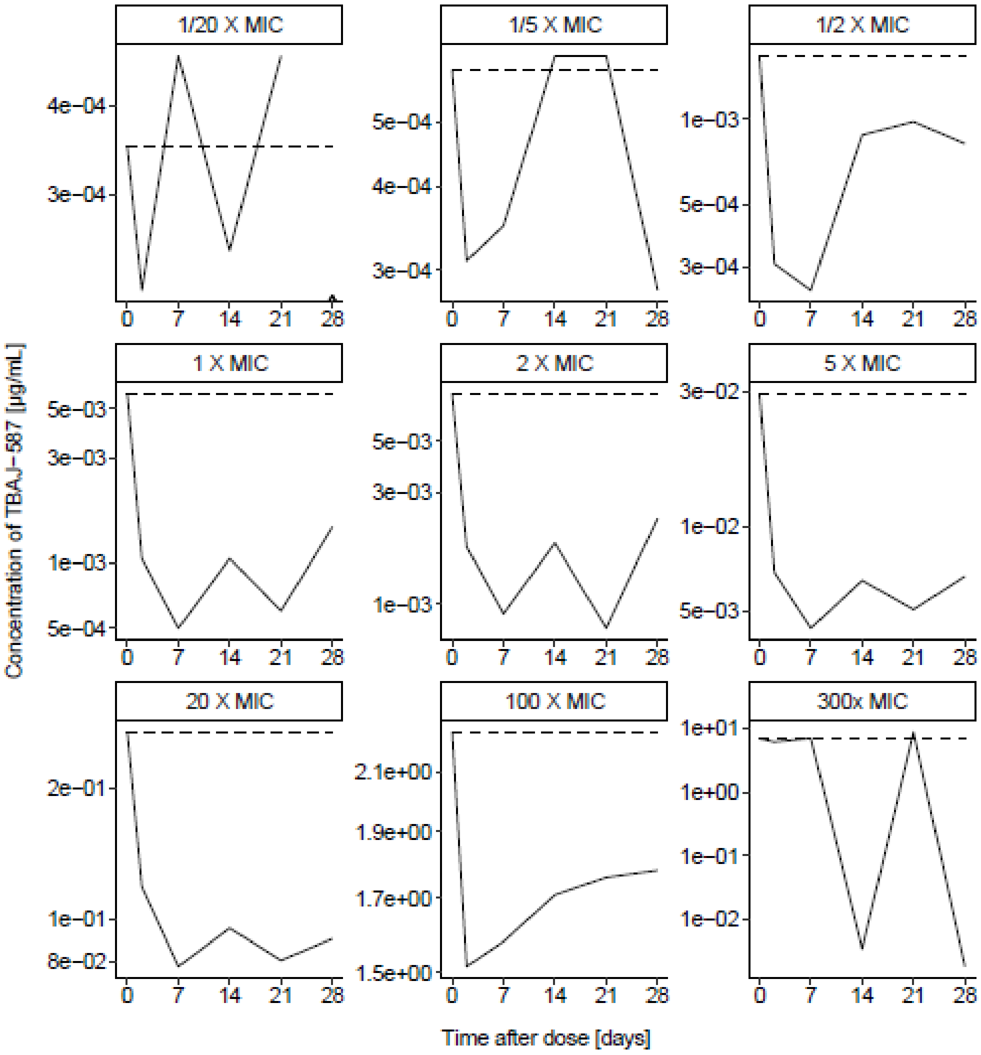

**Figure S4. Drug concentration of M2 in standard broth.** Relative concentration to time zero of M2 in standard medium for the time kill assay, dotted line marks the baseline concentration. Below quantification limit marked on x-axis.

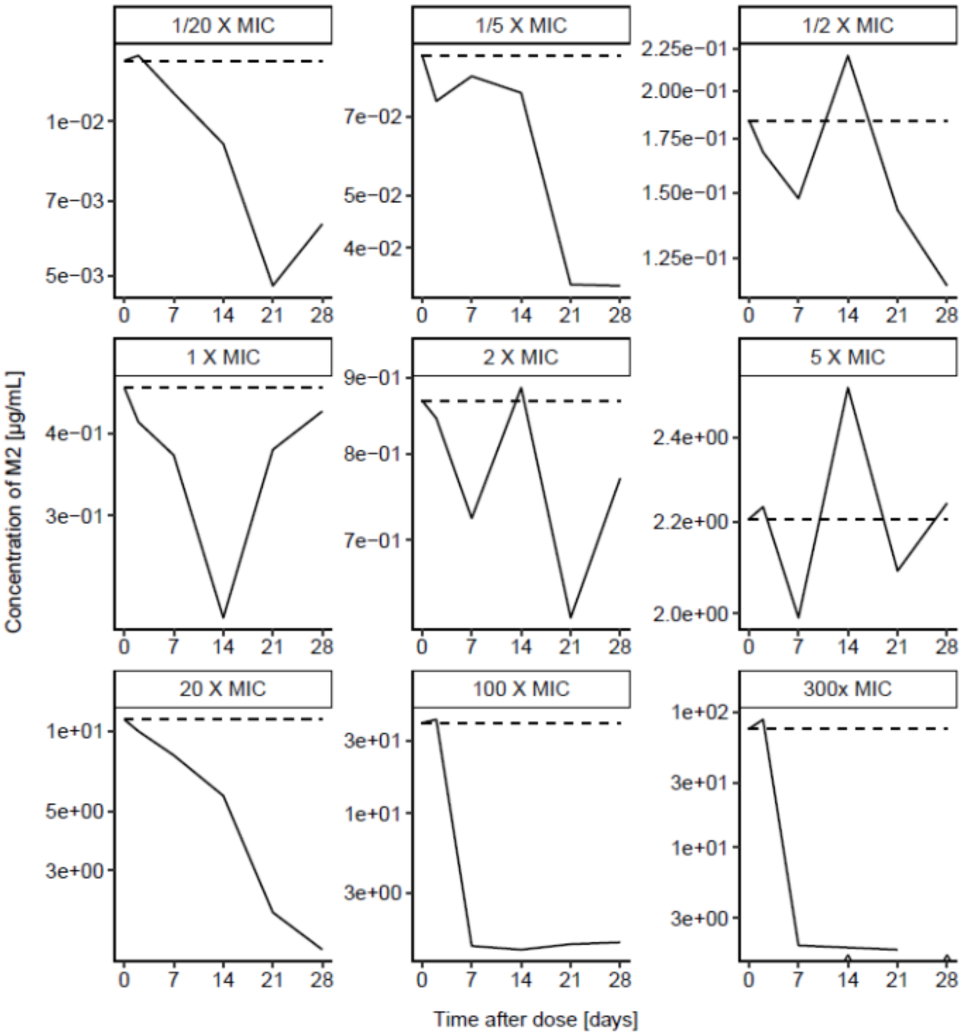

**Figure S5. Drug concentration of M2 in cholesterol broth.** Relative concentration to time zero of M2 in cholesterol medium for the time kill assay, dotted line marks the baseline concentration. Below quantification limit marked on x-axis.

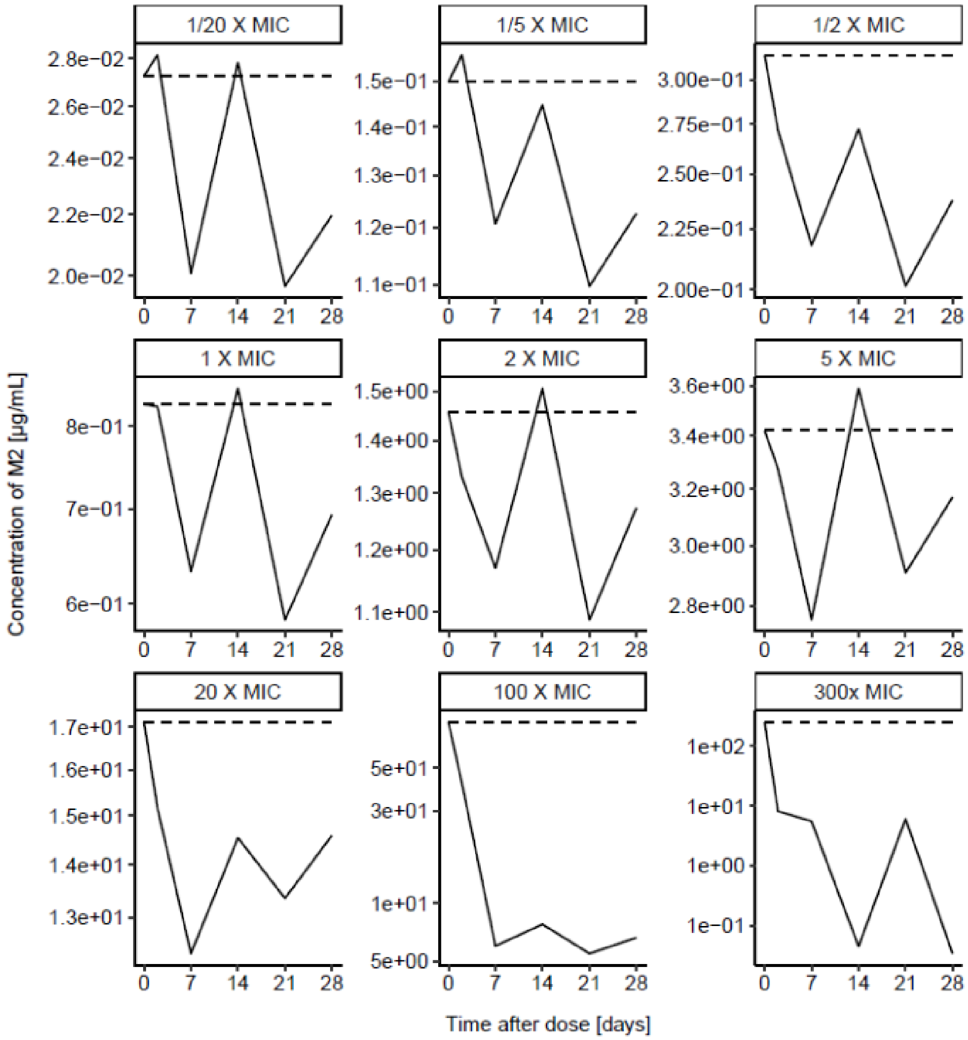

**Figure S6. Drug concentration of M2 in fatty acids broth.** Relative concentration to time zero of M2 in fatty acids medium for the time kill assay, dotted line marks the baseline concentration. Below quantification limit marked on x-axis.

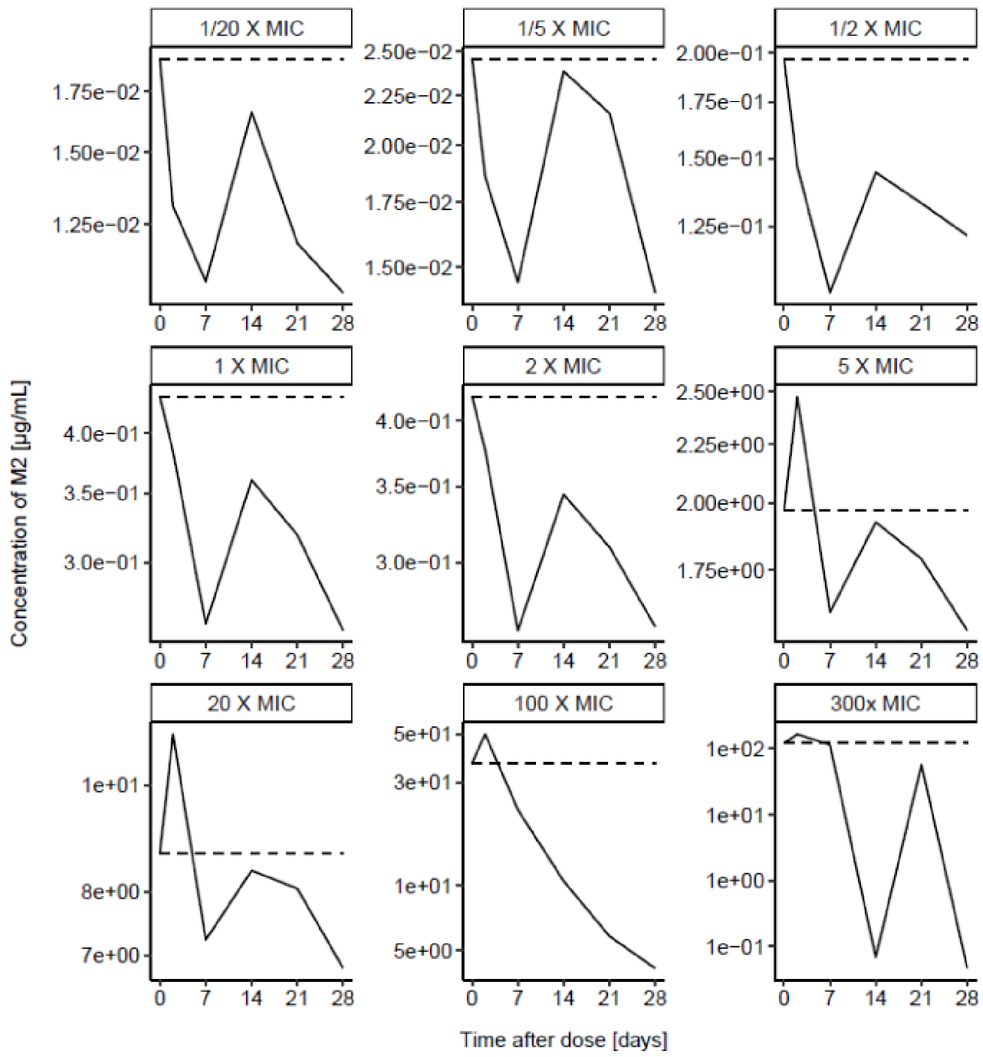

**Figure S7. Drug concentration of M3 in standard broth.** Relative concentration to time zero of M3 in standard medium for the time kill assay, dotted line marks the baseline concentration. Below quantification limit marked on x-axis.

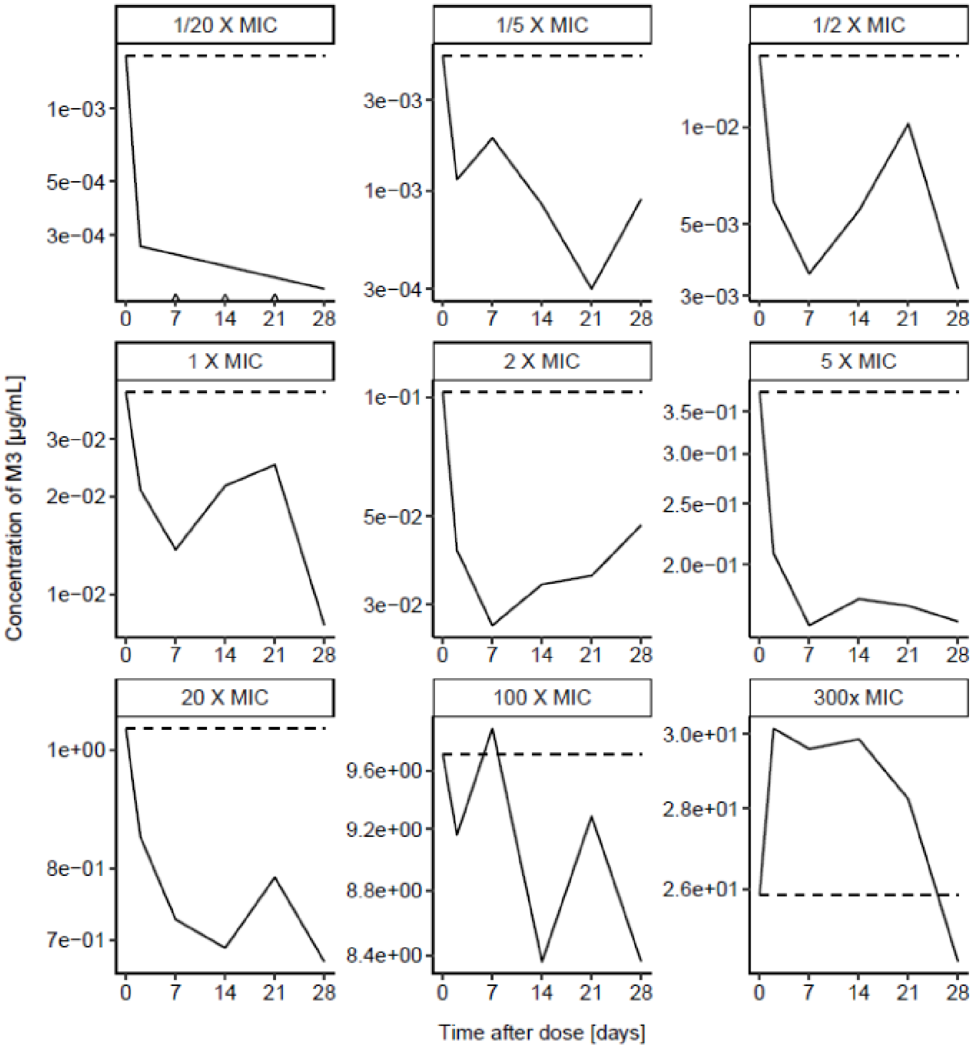

**Figure S8. Drug concentration of M3 in cholesterol broth.** Relative concentration to time zero of M3 in cholesterol medium for the time kill assay, dotted line marks the baseline concentration. Below quantification limit marked on x-axis.

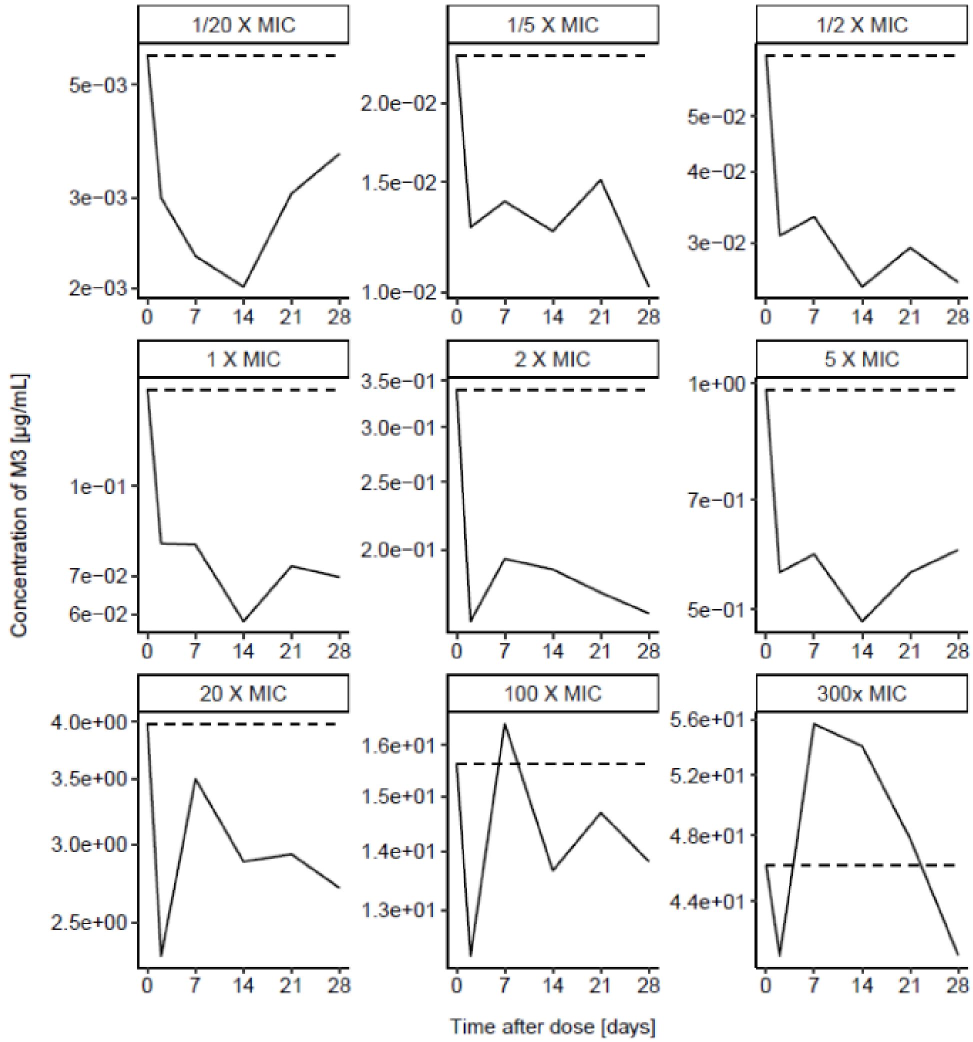

**Figure S9. Drug concentration of M3 in fatty acids broth.** Relative concentration to time zero of M3 in fatty acids medium for the time kill assay, dotted line marks the baseline concentration. Below quantification limit marked on x-axis.

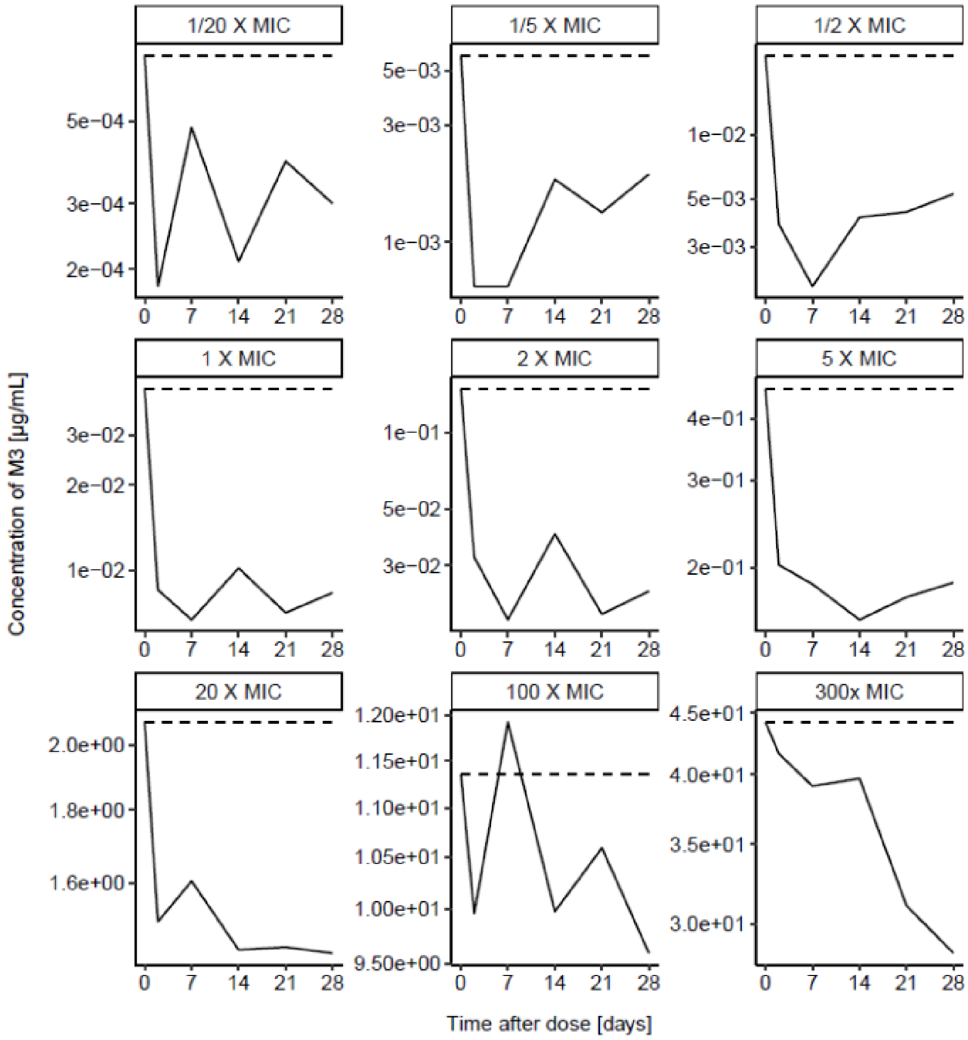

**Figure S10. Drug concentration of M12 in standard broth.** Relative concentration to time zero of M12 in standard medium for the time kill assay, dotted line marks the baseline concentration. Below quantification limit marked on x-axis.

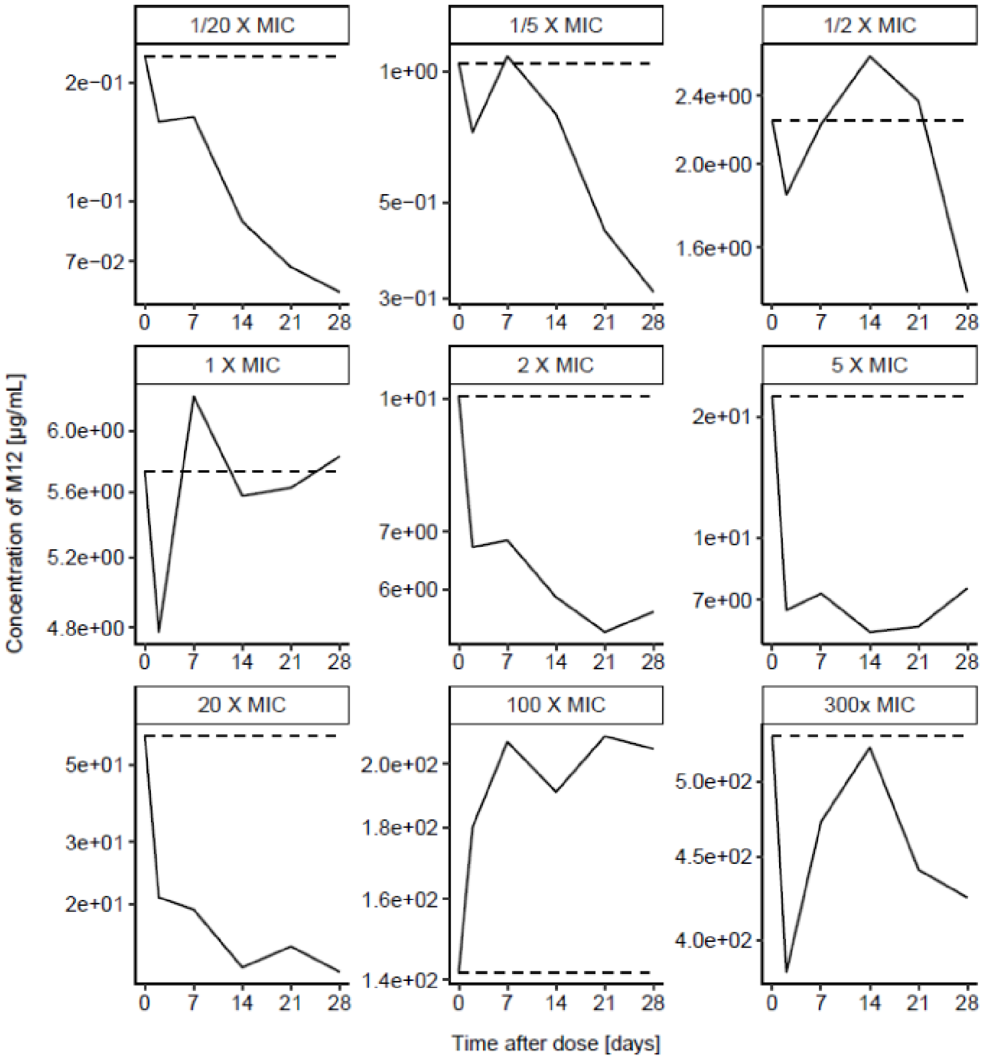

**Figure S11. Drug concentration of M12 in cholesterol broth.** Relative concentration to time zero of M12 in cholesterol medium for the time kill assay, dotted line marks the baseline concentration. Below quantification limit marked on x-axis.

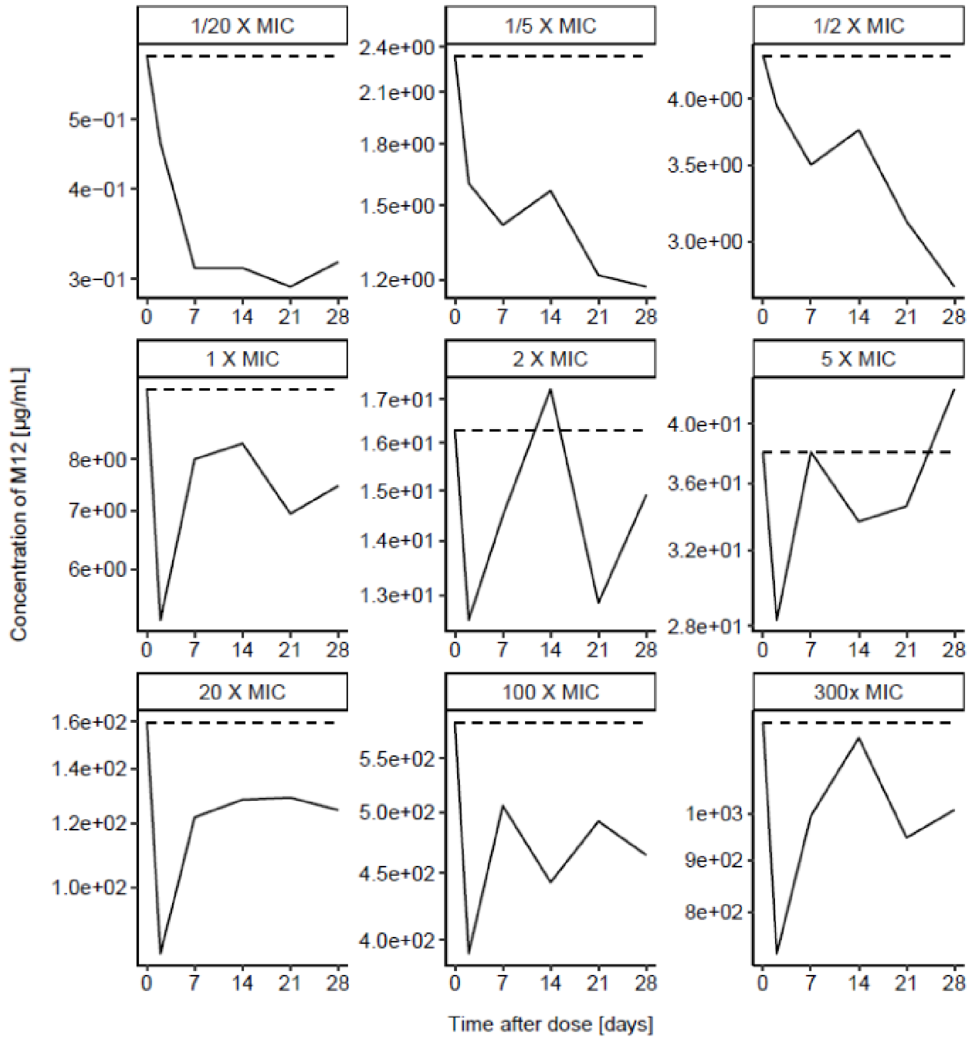

**Figure S12. Drug concentration of M12 in fatty acids broth.** Relative concentration to time zero of M12 in fatty acids medium for the time kill assay, dotted line marks the baseline concentration. Below quantification limit marked on x-axis.

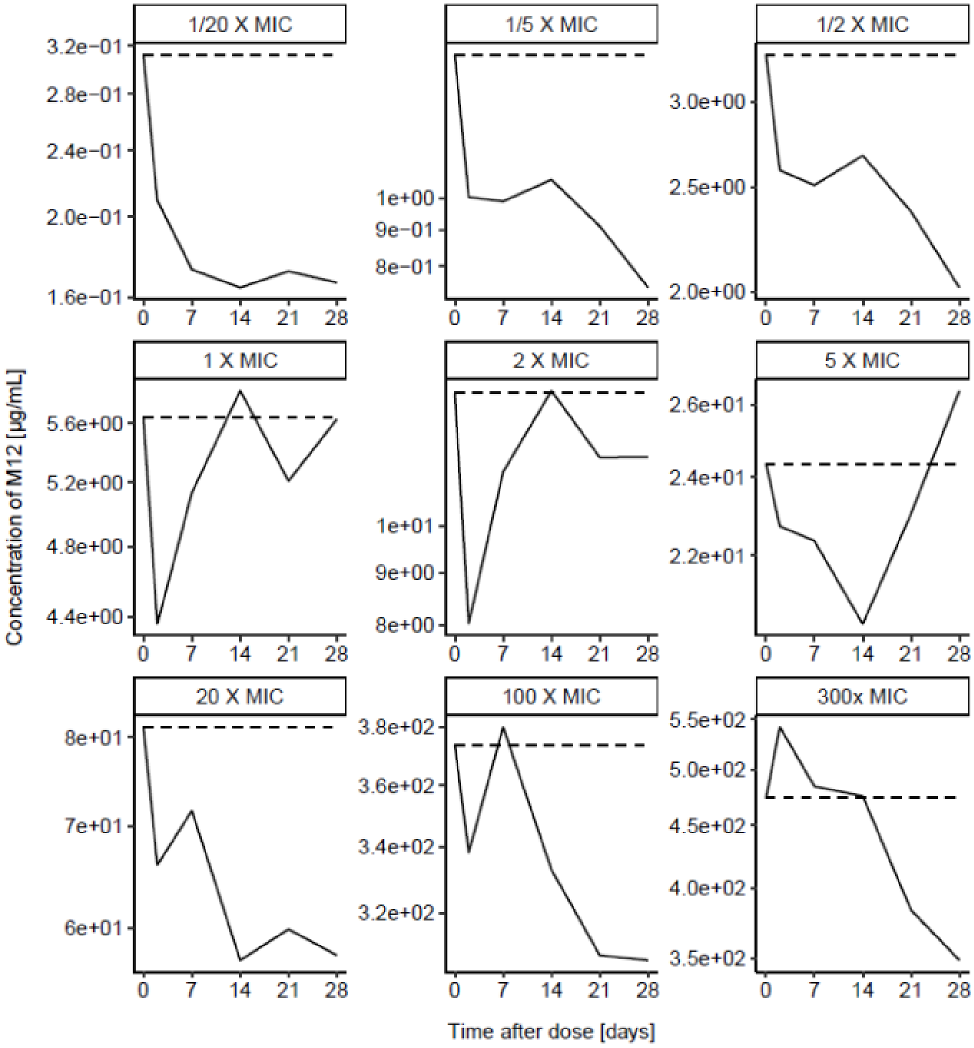

**Figure S13. Stability of TBAJ-587 M3 metabolite.** The recovery of TBAJ-587-M3 metabolite was evaluated in standard 7H9 + 0.5%BSA medium at 10x MIC, without bacteria, in glass and low binding polypropylene tubes over six day at 37°C.

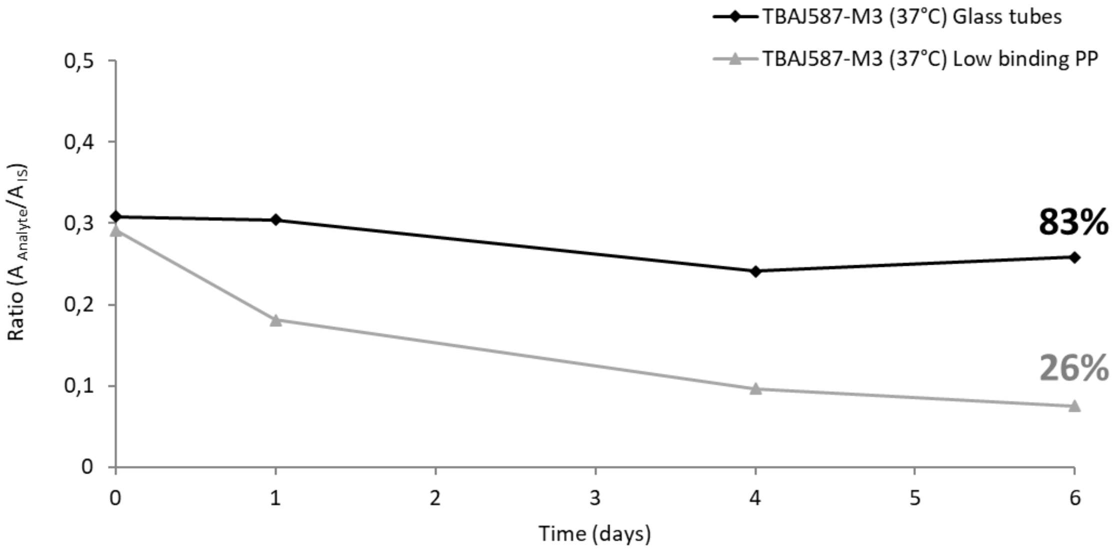
